# JAZF1-SUZ12 dysregulates PRC2 function and gene expression during cell differentiation

**DOI:** 10.1101/2021.04.15.440052

**Authors:** Manuel Tavares, Garima Khandelwal, Joanne Mutter, Keijo Viiri, Manuel Beltran, Jan J. Brosens, Richard G. Jenner

## Abstract

Polycomb repressive complex 2 (PRC2) methylates histone H3 lysine 27 (H3K27me3) to maintain repression of genes specific for other cell types and is essential for cell differentiation. In endometrial stromal sarcoma, the PRC2 subunit SUZ12 is often fused with the NuA4/TIP60 subunit JAZF1. Here, we show that JAZF1-SUZ12 dysregulates PRC2 composition, recruitment, histone modification, gene expression and cell differentiation. The loss of the SUZ12 N-terminus in the fusion protein disrupted interaction with the PRC2 accessory factors JARID2, EPOP and PALI1 and prevented recruitment of PRC2 from RNA to chromatin. In undifferentiated cells, JAZF1-SUZ12 occupied PRC2 target genes but gained a JAZF1-like binding profile during cell differentiation. JAZF1-SUZ12 reduced H3K27me3 and increased H4Kac at PRC2 target genes, and this was associated with disruption in gene expression and cell differentiation programs. These results reveal the defects in chromatin regulation caused by JAZF1-SUZ12, which may underlie its role in oncogenesis.

## INTRODUCTION

Endometrial stromal tumours are a set of uterine malignancies that are divided into four subtypes; benign endometrial stromal nodules (ESN), low-grade endometrial stromal sarcoma (LG-ESS), high-grade endometrial stromal sarcoma (HG-ESS) and undifferentiated uterine sarcoma (UUS). Around 50% of ESN and LG-ESS cases exhibit the chromosomal rearrangement t(7;17)(p15:q21) (Chiang et al., 2011; Micci et al., 2016), which results in production of a JAZF1-SUZ12 fusion protein that comprises the first 128 amino acids of JAZF1 in place of the first 93 amino acids of SUZ12 (Koontz et al., 2001). The WT SUZ12 allele is also silenced in these LG-ESS cases (Li et al 2007). 40% of LG-ESS patients do not respond to current therapeutic regimens, highlighting the need for greater understanding of JAZF1-SUZ12 function and the development of new treatment strategies for this disease (Amant et al., 2007; Beck et al., 2012).

SUZ12 is a core subunit of polycomb repressive complex 2 (PRC2), along with EZH2 or EZH1, EED and RBBP4 or RBBP7 (Margueron and Reinberg, 2011). PRC2 associates with genes encoding developmental regulators specific for other cell types or cell differentiation stages (Azuara et al., 2006; Boyer et al., 2006; Bracken et al., 2006; Lee et al., 2006). At these genes, PRC2 trimethylates histone H3 lysine 27 (H3K27me3), which allows binding of the canonical form of PRC1 and formation of a repressive chromatin structure that maintains genes in a repressed state. Mice lacking *Suz12* die during embryogenesis due to defects in gastrulation (Pasini et al., 2004; Lee et al., 2006), phenocopying *Ezh2* or *Eed*-null mutants (Faust et al, 1995; O’Carroll et al, 2001). This role for SUZ12 in early development is reflected by its requirement for embryonic stem cell (ESC) differentiation *in vitro* (Lee et al., 2006; Pasini et al., 2007; Riising et al., 2014). PRC2 is also required for the differentiation of numerous other cell types, both during embryogenesis and throughout life, and is frequently dysregulated in cancer (Comet et al., 2016; Deevy & Bracken, 2019; Prezioso & Orlando, 2011; Schuettengruber et al., 2017).

In addition to the core PRC2 subunits, a number of accessory factors have been discovered that define two variants of PRC2; PRC2.1 and PRC2.2 (Alekseyenko et al, 2014; Beringer et al., 2016; Conway et al., 2018; Grijzenhout et al., 2016; Hauri et al., 2016; Liefke et al, 2016). PRC2.1 is composed of the PRC2 core plus one of PCL1, PCL2 or PCL3 (also named PHF1, MTF2 and PHF19) along with either EPOP (Alekseyenko et al., 2014; Beringer et al., 2016; Liefke et al., 2016; Zhang et al., 2011), PALI1 or PALI2 (Alekseyenko et al., 2014; Conway et al., 2018; Zhang et al., 2011). PRC2.2 is less complex and is formed by the PRC2 core plus AEBP2 and JARID2 (Alekseyenko et al., 2014; Grijzenhout et al., 2016; Hauri et al., 2016). The accessory subunits act in combination to recruit PRC2 to its target sites (Højfeldt et al., 2018; Højfeldt et al., 2019; Oksuz et al., 2018; Perino et al., 2018; Wang et al., 2017a; Youmans et al., 2018) and are essential for normal development (Brien et al., 2012; Conway et al., 2018; Grijzenhout et al., 2016; Landeira et al., 2010; Li et al., 2010, 2017; Liefke et al., 2016; Shen et al., 2009; Walker et al., 2010; Zhang et al., 2011). SUZ12 interacts with EZH2 and EED through its C-terminal VEFS domain, and with JARID2, AEBP2, EPOP and PCL proteins through the N-terminal part of the protein (Chen et al., 2018; Kasinath et al., 2018; Kloet et al., 2016; Youmans et al., 2018). Recently, an additional factor, CXorf67/EZHIP/CATACOMB, was discovered to interact with core PRC2 and inhibit its histone methyltransferase activity (Hübner et al., 2019; Jain et al., 2019; Piunti et al., 2019; Ragazzini et al., 2019).

In addition to binding to chromatin, PRC2 also interacts with RNA. First found to bind specific long non-coding RNAs, PRC2 has since been found to primarily interact with nascent pre-mRNA in cells (Beltran et al., 2016; Davidovich et al., 2013; Hendrickson et al., 2016; Kaneko et al., 2013; Zhao et al., 2010), with a preference for G-quadruplex-forming G-tract repeats (Beltran et al., 2019; Wang et al., 2017b). *In vitro*, G-quadruplex RNA inhibits the interaction of PRC2 with DNA, nucleosomes and H3 tails (Beltran et al., 2016, Beltran et al., 2019; Long et al., 2017; Wang et al., 2017a; Zhang et al., 2019) and, in ESC, transcriptional inhibition (Riising et al., 2014), polyA-site insertion (Kaneko et al., 2014), and RNA degradation (Beltran et al., 2016) induce PRC2 recruitment to active genes. Reciprocally, blocking nuclear RNA degradation (Garland et al., 2019) or tethering G-tract RNAs to polycomb target genes removes PRC2 from chromatin (Beltran et al., 2019), supporting a model in which nascent RNA restricts PRC2 recruitment to genes that are already silent (Kaneko et al., 2014, Beltran et al., 2016; Comet et al., 2016; Hosogane et al., 2016; Riising et al., 2014).

The effect of JAZF1-SUZ12 on PRC2 function is not well understood. Ectopic expression of JAZF1-SUZ12 and knockdown of endogenous SUZ12 in HEK293 cells increased cell proliferation and resistance to hypoxia (Li et al., 2007). JAZF1 fusion has been reported to reduce interaction of SUZ12 with EZH2 and EED and consequently reduce PRC2 H3K27 methyltransferase activity (Ma et al., 2016). JAZF1-SUZ12 has also been found to lack interaction with JARID2 and EPOP (Chen et al., 2018) but its interaction with other PRC2 accessory factors has not been measured.

Compared to SUZ12, understanding of JAZF1 function has been limited. JAZF1 interacts with the orphan nuclear receptor NR2C2 (TAK1/TR4), inhibiting its activity (Nakajima et al., 2004) and regulates genes with functions in translation and RNA splicing (Kobiita et al., 2020; Procida et al., 2021). Shedding more light on the mechanism of JAZF1 action, the protein was recently found to be associated with the NuA4/TIP60 histone acetyltransferase complex (Piunti et al., 2019; Procida et al., 2021) which exchanges H2A for H2A.Z (Nishibuchi et al., 2014; Xu et al., 2012) and acetylates histone H4 (H4Kac), as well as histone H2A and its variants (Steunou et al., 2014). Correspondingly, fusion of JAZF1-SUZ12 results in ectopic interaction between PRC2 and NuA4 components (Piunti et al., 2019). The association of JAZF1 with NuA4 is consistent with reports of other fusion events between PRC2 and NuA4 subunits in LG-ESS, namely EPC1-PHF1 (Micci et al., 2006), JAZF1-PHF1 (Panagopoulos et al., 2008), MEAF6-PHF1(Panagopoulos et al., 2012), MBTD1-CXorf67/EZHIP (Dewaele et al., 2014), BRD8-PHF1(Davidson and Micci, 2017), MEAF6-SUZ12 (Makise et al., 2019) and MBTD1-PHF1 (Han et al., 2020), and indicates that the fusion of PRC2 with NuA4 drives oncogenesis in all of these cases. However, how JAZF1-SUZ12 affects PRC2 composition, recruitment and patterns of H3K27me3 and the impact of these changes on gene expression and cell differentiation remain unclear.

Here, we show that JAZF1-SUZ12 disrupts interaction of PRC2 with JARID2, EPOP and PALI1, and dysregulates PRC2 recruitment to chromatin. We also separate changes due to loss of the SUZ12 N-terminus from those caused by fusion to JAZF1. JAZF1-SUZ12 chromatin occupancy diverges from WT SUZ12 during cell differentiation and resulted in reduced H3K27me3 and increased H4Kac at PRC2 target genes. ESC expressing JAZF1-SUZ12 show altered patterns of gene expression and disrupted embryoid body (EB) formation and JAZF1-SUZ12 also drives gene expression changes in primary human endometrial stromal cells (hEnSC). We propose that these effects contribute to JAZF1-SUZ12-driven oncogenesis.

## RESULTS

### JAZF1-SUZ12 lacks interaction with JARID2, EPOP and PALI1 due to loss of the SUZ12 N-terminus

We first sought to establish the effect of JAZF1 fusion with SUZ12 on PRC2 composition. To address this, we generated *Suz12*^GT/GT^ ESC lines stably expressing FLAG-tagged SUZ12, JAZF1-SUZ12, JAZF1 or GFP (Figure 1A). To distinguish the changes in PRC2 composition due to loss the of the N-terminal 93 amino acids of SUZ12 and the changes due to gain of the first 128 amino acids of JAZF1, we also generated a cell line stably expressing FLAG-tagged SUZ12Δ93. Immunoblotting confirmed that the levels of SUZ12, SUZ12Δ93 and JAZF1-SUZ12 proteins were similar to each other and to endogenous SUZ12 (Figure S1A).

**Figure 1.**
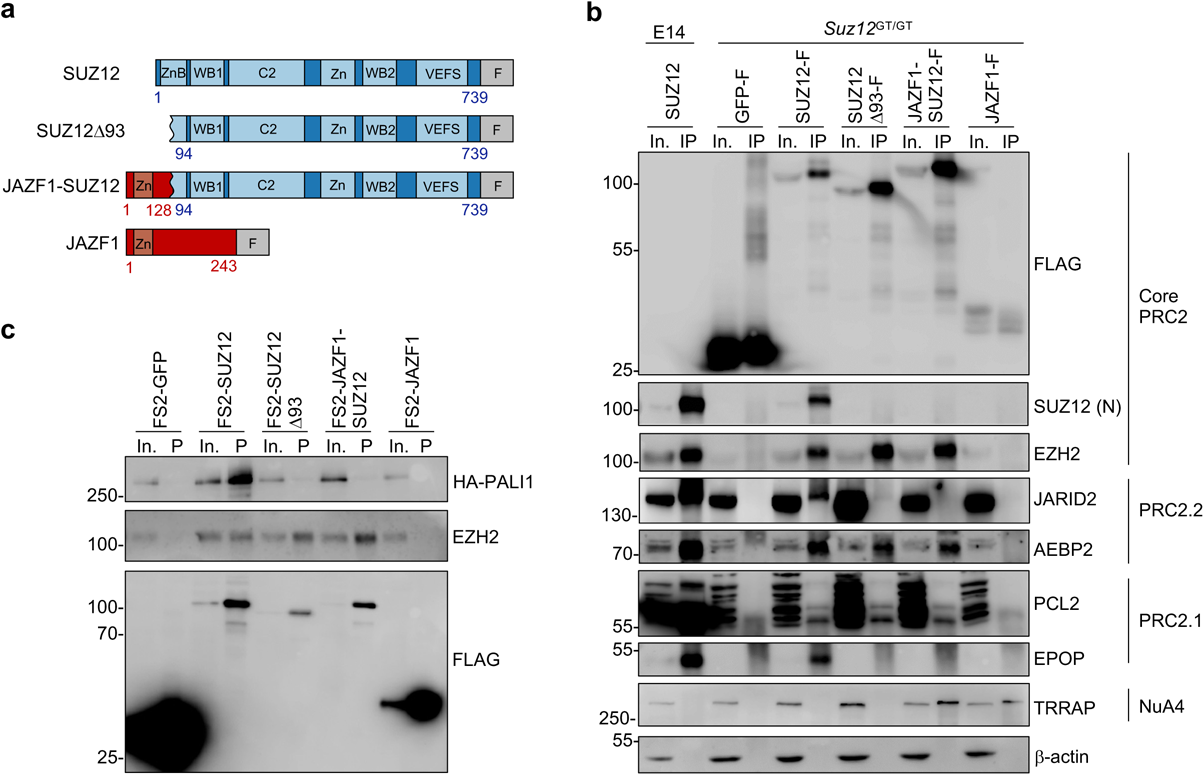
JAZF1-SUZ12 lacks interaction with JARID2, EPOP and PALI1 due to loss of the SUZ12 N-terminus. **a.** Models of SUZ12, SUZ12Δ93, JAZF1-SUZ12 and JAZF1 constructs with C-terminal FLAG tags (F). **b.** Immunoblots for FLAG, SUZ12 (N-terminus), EZH2, JARID2, AEBP2, PCL2, EPOP, TRRAP and β-actin in input (In.) and SUZ12 IPs from WT E14 cells or input and FLAG IPs from *Suz12*^GT/GT^ cells stably expressing FLAG-tagged GFP, SUZ12, SUZ12Δ93, JAZF1-SUZ12 or JAZF1. **c.** Immunoblots for FLAG, HA-PALI and EZH2 in input and Strep-Tactin pull-downs (P) from NIH-3T3 cells transfected with HA-tagged PALI1 and FS2-tagged GFP, SUZ12, SUZ12Δ93, JAZF1-SUZ12 or JAZF1. **d.** See also Figure S1.

We then measured the effect of SUZ12Δ93 and JAZF1-SUZ12 on PRC2 composition by co-immunoprecipitation and immunoblotting (Figure 1B). Co-immunoprecipitation of endogenous SUZ12 from WT ESC provided an additional control. This revealed that SUZ12, SUZ12Δ93 and JAZF1-SUZ12 interacted with the core subunit EZH2 and the accessory subunits AEBP2 and PCL2. However, unlike SUZ12, SUZ12Δ93 and JAZF1-SUZ12 did not interact with JARID2 or EPOP. These findings were confirmed by reciprocal co-IPs of tagged forms of AEBP2, EPOP and PCL3 (Figure S1B-D).

To determine the effect of JAZF1-SUZ12 on the association of PALI1 with PRC2, we co-transfected FS2-tagged SUZ12, SUZ12Δ93, JAZF1-SUZ12 and JAZF1 constructs with HA/FLAG-tagged PALI1 and performed Strep-Tactin affinity purification (Figure 1C). We found that PALI1 did not interact with SUZ12Δ93 or JAZF1-SUZ12, thus following the same pattern as EPOP and JARID2.

We also sought to confirm recent findings that JAZF1-SUZ12 exhibited an ectopic interaction with the NuA4/TIP60 complex (Piunti et al., 2019) We found that IP for JAZF1-SUZ12 or JAZF1 co-precipitated the NuA4 subunit TRRAP but that this wasn’t the case for IP of SUZ12 or SUZ12Δ93 (Figure 1B). Therefore, we conclude that fusion of SUZ12 with JAZF1 disrupts interaction with EPOP, JARID2 and PALI1 due to loss of the SUZ12 N-terminus and induces interaction of SUZ12 with TRRAP due to gain of the N-terminus of JAZF1.

### Lack of recruitment of JAZF1-SUZ12 to chromatin upon RNA depletion due to loss of interaction with JARID2 and EPOP

The recruitment of PRC2 to chromatin is antagonised by RNA and this can be observed by the increase in PRC2 chromatin association upon RNA degradation in cells (Beltran et al., 2016, Beltran et al., 2019). Thus, we sought to determine the effect of JAZF1-SUZ12 on the recruitment of PRC2 to chromatin upon RNaseA treatment. ESC expressing SUZ12, SUZ12Δ93 or JAZF1-SUZ12 were permeabilised, mock-treated or treated with RNaseA and the nucleoplasmic and chromatin fractions purified. This confirmed that RNaseA treatment increased the association of WT SUZ12 with chromatin but decreased the association of FUS with chromatin (Figure 2A), as previously described (Beltran et al., 2016). In contrast, SUZ12Δ93 and JAZF1-SUZ12 were both depleted from chromatin by RNaseA treatment, as was EZH2 in cells expressing these proteins. Thus, loss of the SUZ12 N-terminus prevents the recruitment of JAZF1-SUZ12 from RNA to chromatin.

**Figure 2.**
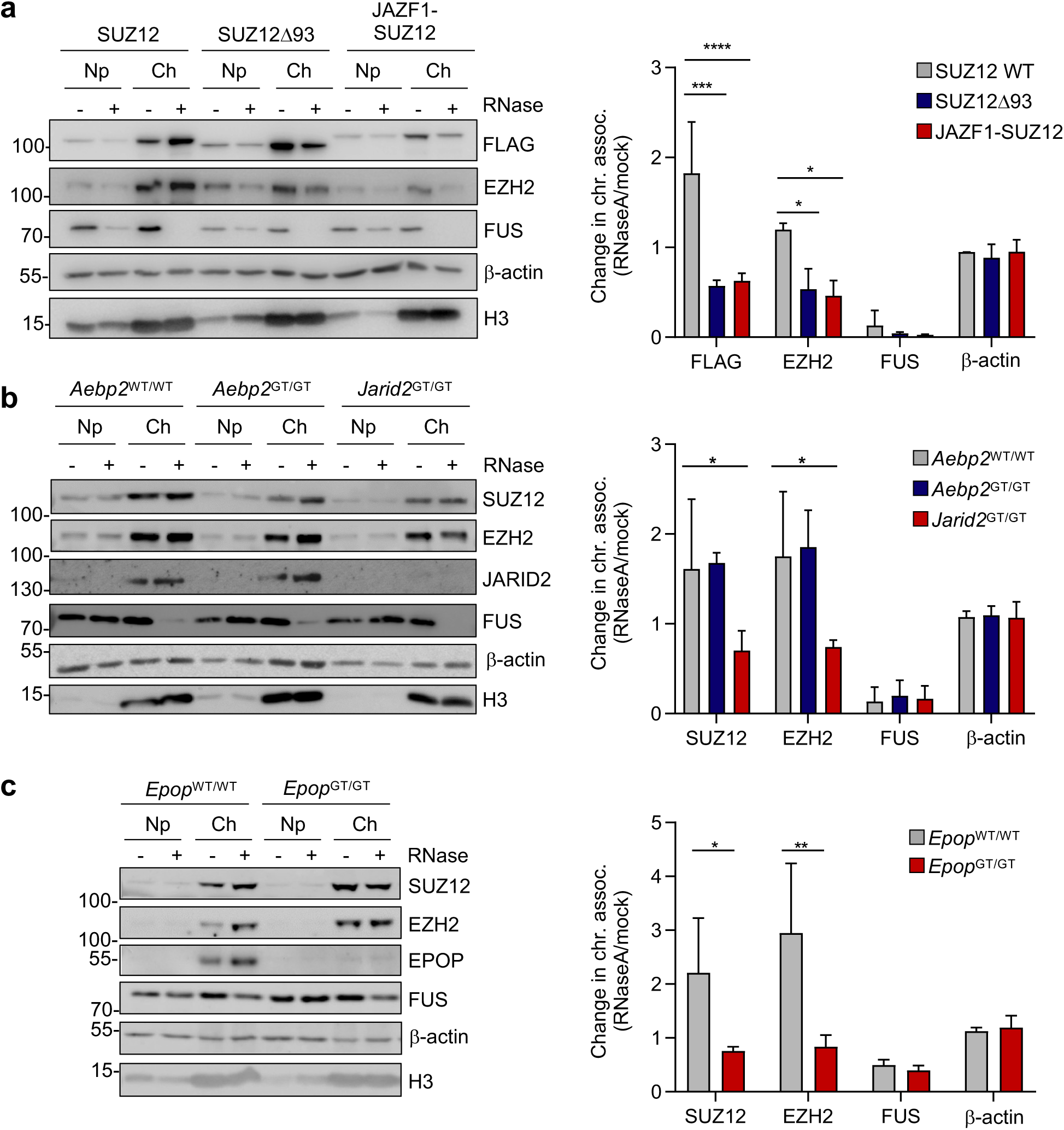
JAZF1-SUZ12 is not recruited to chromatin upon RNA depletion due to loss of interaction with JARID2 and EPOP. **a.** *Left:* Immunoblots for FLAG, EZH2, FUS, β-actin and H3 in nucleoplasm (Np) and chromatin (Ch) fractions from mock or RNaseA-treated *Suz12*^GT/GT^ cells expressing either FLAG-tagged SUZ12, SUZ12Δ93 or JAZF1-SUZ12. Representative of three independent experiments. *Right*: Fold change in FLAG, EZH2, FUS and β-actin in the chromatin fraction upon RNaseA treatment (mean and s.d, n=3). **b.** As a., but for endogenous SUZ12, EZH2, JARID2, FUS, β-actin and H3 in *Aebp2*^WT/WT^*, Aebp2*^GT/GT^ *and Jarid2*^GT/GT^ ESC. **c.** As a., but for endogenous SUZ12, EZH2, EPOP, FUS, β-actin and H3 in *Epop*^WT/WT^ and *Epop*^GT/GT^ ESC. See also Figure S2.

Given that SUZ12Δ93 and JAZF1-SUZ12 lack interaction with JARID2, EPOP and PALI1, we asked whether this was the reason that these proteins were not recruited to chromatin upon RNA degradation. We first tested the effect of loss of the PRC2.2 subunits AEBP2 or JARID2 on PRC2 chromatin association upon RNA degradation. *Aebp2*^WT/WT^, *Aebp2*^GT/GT^ and *Jarid2*^GT/GT^ cells were permeabilized, mock or RNaseA-treated and fractionated as before (Figure 2B). We found that loss of AEBP2 had no effect on the recruitment of SUZ12 and EZH2 to chromatin upon RNA depletion. In contrast, loss of JARID2 abrogated the recruitment of PRC2 subunits to chromatin (Figure 2B). These results show that within PRC2.2, JARID2 is necessary PRC2 recruitment from RNA to chromatin, while AEBP2 is not.

We next determined the requirement for the PRC2.1 subunits EPOP, PALI1 and PCL1-3 on the recruitment of PRC2 to chromatin upon RNA depletion. In cells lacking EPOP, we found there was no enrichment of SUZ12 and EZH2 in the chromatin fraction upon RNaseA treatment (Figure 2C). In contrast, we did not observe a change in the recruitment of SUZ12 to chromatin in cells lacking PALI1 or PCL1-3 (Figures S2A and S2B). We conclude that JAZF1-SUZ12-containing PRC2 is not recruited from RNA to chromatin and that this is due to lack of interaction with JARID2 and EPOP.

### JAZF1-SUZ12 takes on a JAZF1-like binding profile during cell differentiation

Considering that JAZF1-SUZ12 displayed defects in recruitment from RNA to chromatin, we next asked how fusion to JAZF1 affected the pattern of SUZ12 chromatin occupancy during cell differentiation. To measure this, we performed calibrated ChIP-seq for FLAG-tagged GFP, SUZ12, SUZ12Δ93, JAZF1-SUZ12 and JAZF1 in ESC and at 4 and 8 days after induction of EB formation. We first identified binding sites for each protein at each timepoint and used hierarchical clustering to determine how these were related to one another (Figure 3A). This showed that SUZ12 and JAZF1 samples clustered separately, demonstrating they had distinct binding profiles. SUZ12Δ93 samples clustered together with SUZ12 indicating that loss of SUZ12 N-terminus did not completely disrupt the pattern of PRC2 chromatin occupancy. Strikingly, the JAZF1-SUZ12 binding profile resembled SUZ12 and SUZ12Δ93 in ESC but in EBs, its binding profile was more similar to that of JAZF1. This shift of JAZF1-SUZ12 towards a JAZF1 binding profile during ESC differentiation was also apparent by plotting SUZ12 and JAZF1 occupancy at JAZF1-SUZ12 binding sites in ESC compared with JAZF1-SUZ12 binding sites in EBs (Figure 3B). Indeed, quantifying the number of binding sites shared between the different factors revealed that JAZF1-SUZ12 occupied 39% of JAZF1 binding sites in ESC, which increased to 82% of JAZF1 binding sites in day 8 EBs (Figure S3A). Our analysis also revealed that JAZF1-SUZ12 occupied a large number of sites that were not shared with SUZ12 or JAZF1 and that the number of these ectopic binding sites also increased during ESC differentiation (Figure S3A).

**Figure 3.**
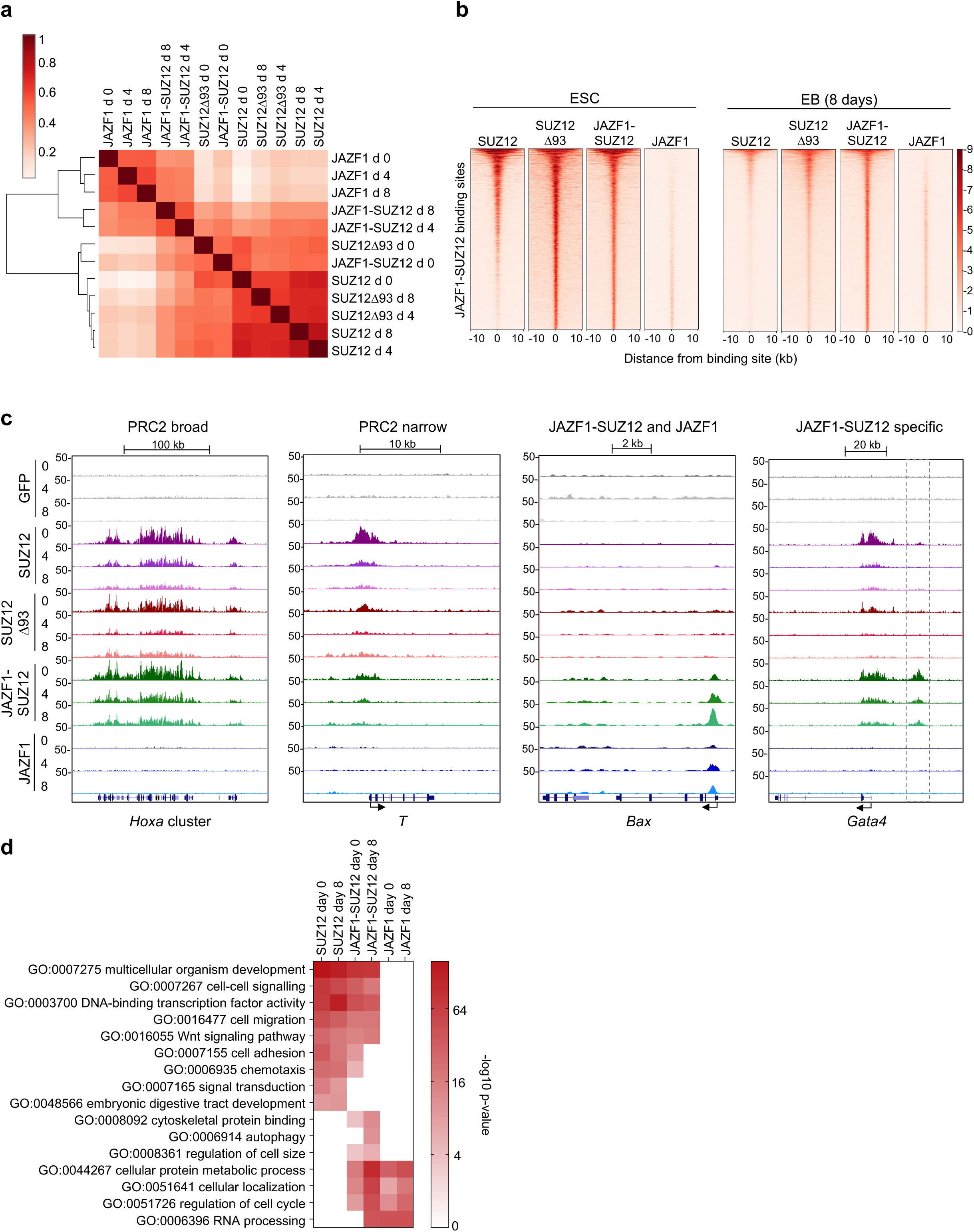
JAZF1-SUZ12 takes on a JAZF1-like binding profile during cell differentiation. **a.** Hierarchical clustering showing the relationship between the patterns of SUZ12, SUZ12Δ93, JAZF1-SUZ12 and JAZF1 genome occupancy in ESC and in embryoid bodies (EB) 4 and 8 days after induction of EB formation. Pearson correlations between samples are shown by colour, according to the scale on the left. **b.** Heatmaps showing SUZ12, SUZ12Δ93, JAZF1-SUZ12 and JAZF1 occupancy at sites bound by JAZF1-SUZ12 in ESC (day 0) and sites bound by JAZF1-SUZ12 in EBs (day 8). Occupancy (normalised reads) is indicated by color, according to the scale on the right. At each timepoint, sites are ordered by the average occupancy of all factors, from high to low. **c.** Four patterns of SUZ12, SUZ12Δ93, JAZF1-SUZ12 and JAZF1 genome occupancy (normalised reads per 10 bp window) in ESC at 0, 4 and 8 days after initiation of differentiation into EBs. Left: a broad region of shared SUZ12, SUZ12Δ93 and JAZF1-SUZ12 occupancy at the *Hoxa* cluster. Center left: a narrow region of SUZ12 occupancy at *T* that exhibits reduced SUZ12Δ93 and JAZF1-SUZ12 binding. Center right: shared JAZF1-SUZ12 and JAZF1 occupancy at *Bax.* Right: JAZF1-SUZ12-specific binding upstream of *Gata4* not shared by SUZ12 or JAZF1 (dashed box). **d.** Heatmap showing enrichment (-log_10_ p-value) of representative GO terms within the sets of genes occupied by SUZ12, JAZF1-SUZ12 or JAZF1 in ESC or EBs (day 8). See also Figure S3.

We also assessed how the changes in SUZ12 genome occupancy that occur during differentiation were affected by fusion to JAZF1. Comparing SUZ12 and SUZ12Δ93 occupancy between ESC and EBs demonstrated that these proteins tended to be depleted from their target genes during differentiation (Figure S3B), as previously observed for endogenous PRC2 (Kloet et al., 2016). In contrast, JAZF1-SUZ12 tended to be retained at its target genes during ESC differentiation.

These differences JAZF1-SUZ12 and SUZ12 occupancy could also be observed at individual genes (Figures 3C and S3C). At genes with broad regions of SUZ12 binding, for example the *Hoxa* cluster, *Foxa1* or *Pax3*, SUZ12 and JAZF1-SUZ12 occupancy was similar, with extensive binding in ESC that decreased during differentiation. At genes with more narrow regions of SUZ12 occupancy, for example *T* (Brachyury), *Bmp6* or *Fgf5*, JAZF1-SUZ12 binding was greatly reduced. This was also the case for SUZ12Δ93, indicating that the reduction in JAZF1-SUZ12 binding at these sites was due to lack of the SUZ12 N-terminus. In contrast, JAZF1-SUZ12 binding could be observed at other genes, such as *Bax, Cbx3* and *Mcl1*, that lacked SUZ12 occupancy but were instead occupied by JAZF1. At these genes, JAZF1-SUZ12 occupancy increased during ESC differentiation. Finally, sites occupied by JAZF1-SUZ12 but not SUZ12, SUZ12Δ93 or JAZF1 could be identified, for example a region upstream of *Gata4* (Figure 3C), while other sites from which SUZ12 and SUZ12Δ93 were depleted during differentiation instead retained JAZF1-SUZ12, for example *Prdm8* and *Hhex* (Figure S3C).

We conclude from these analyses that loss of the SUZ12 N-terminus reduces JAZF1-SUZ12 recruitment to PRC2 target genes, especially those with narrower regions of occupancy, and that fusion to JAZF1 triggers recruitment to JAZF1 target sites and to an additional set of sites that lack SUZ12 or JAZF1 occupancy.

To determine how these changes in chromatin occupancy might impact cell state, we used Gene Ontology to identify functional terms enriched in the sets of genes associated with SUZ12, JAZF1-SUZ12, or JAZF1 binding sites (Figure 3D and Table S1). We found that some functional annotations, for example those related to development and transcription factor activity, were shared between the sets of genes occupied by SUZ12 and JAZF1-SUZ12, but not by genes occupied by JAZF1. In contrast, other functional gene classes targeted by SUZ12, for example genes with functions in cell adhesion and chemotaxis, were lost from the set of genes occupied by JAZF1-SUZ12, either in both ESC and day 8 EBs or only in EBs. This is consistent with reduced binding of JAZF1-SUZ12 to some SUZ12 target sites (Figures 3C and S3C). Instead of these gene functions, JAZF1-SUZ12 target genes exhibited functions in common with genes bound by JAZF1, including terms related to cell metabolism, cellular localisation and cell cycle, and also functions that were specific to JAZF1-SUZ12 target genes, such as cytoskeletal protein binding and regulation of cell size. Thus, fusion to JAZF1 alters SUZ12 chromatin occupancy and this changes the functional classes of genes targeted by the protein.

### JAZF1-SUZ12 reduces H3K27me3 and increases H4Kac at PRC2 target genes

We next considered the effect of JAZF1-SUZ12 on H3K27me3, catalysed by PRC2, and H4Kac, catalysed by NuA4/TIP60. To do this, we performed calibrated ChIP-seq for H3K27me3 and pan-H4Kac in ESC and EBs expressing WT SUZ12, SUZ12Δ93 or JAZF1-SUZ12. We first examined the effect of JAZF1-SUZ12 on H3K27me3 at canonical PRC2 target sites. To allow measurement of the effect of JAZF1-SUZ12 on H3K27me3 separately from its effect on PRC2 occupancy, we plotted H3K27me3 at binding sites shared by SUZ12, SUZ12Δ93 and JAZF1-SUZ12 (Figure 4A). We found that ESC expressing JAZF1-SUZ12 exhibited lower levels of H3K27me3 at PRC2 target sites than cells expressing WT SUZ12. Examining individual genes revealed a reduction in H3K27me3 even at loci with a broad region of SUZ12 binding at which JAZF1-SUZ12 occupancy was maintained, such as the *Hoxa* cluster, *Foxa1* and *Pax3* (Figures 4B and S4A). At genes with more narrow regions of SUZ12 occupancy from which JAZF1-SUZ12 binding was lost, such as *T*, *Bmp6* and *Fgf5*, H3K27me3 was completely abrogated in ESC expressing JAZF1-SUZ12 (Figures 4B and S4A). This was also observed in cells expressing SUZ12Δ93, indicating that the reduction in H3K27me3 at PRC2 target sites in cells expressing JAZF1-SUZ12 was due to loss of the SUZ12 N-terminus. Thus, we conclude that one way in which fusion of JAZF1 to SUZ12 alters chromatin state is by reducing H3K27me3 at PRC2 target sites.

**Figure 4.**
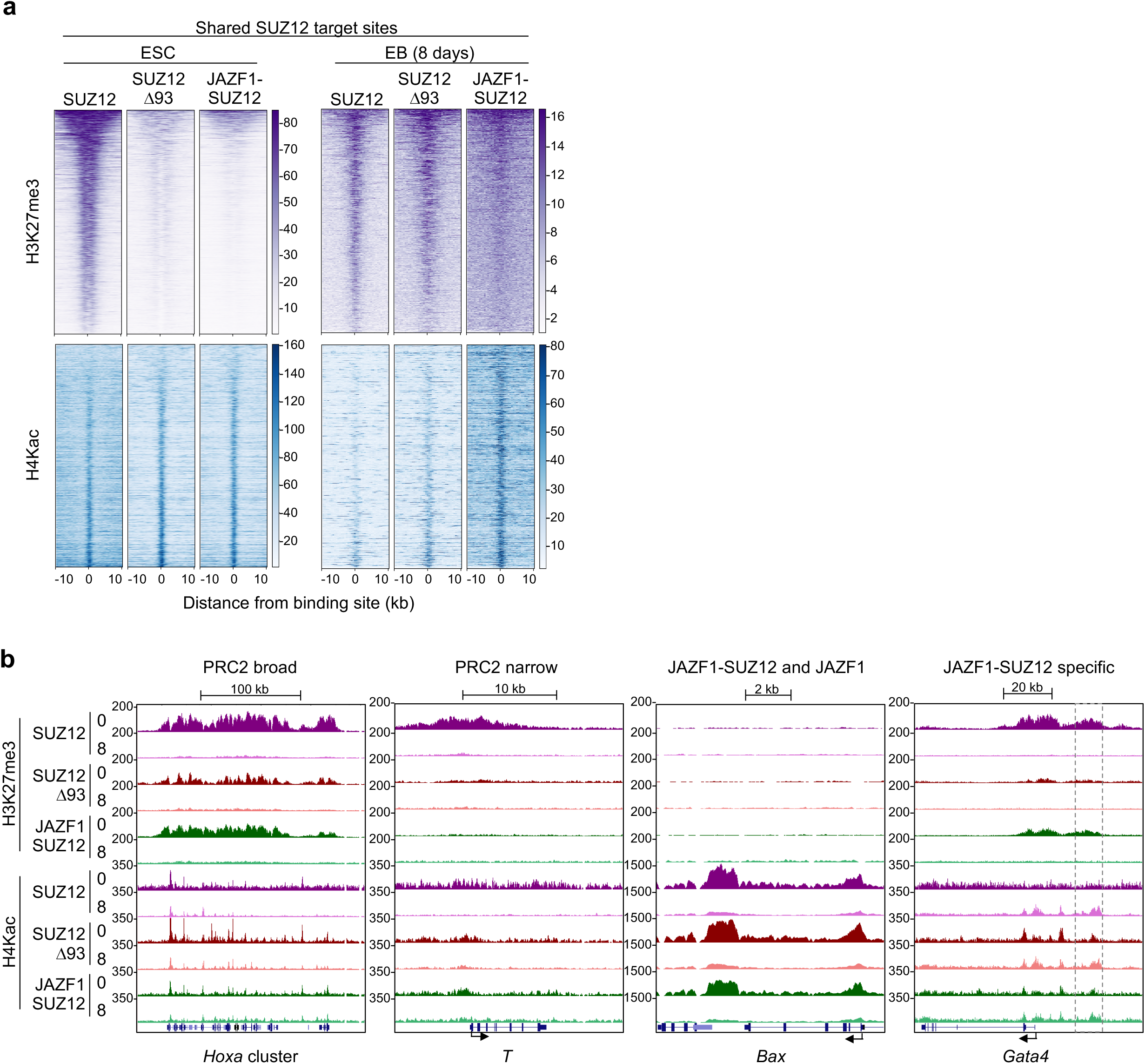
JAZF1-SUZ12 decreases H3K72me3 and increases H4Kac at PRC2 target genes. **a.** Heatmaps of H3K27me3 (purple, top) and H4Kac (blue, bottom) around sites shared by SUZ12, SUZ12Δ93 and JAZF1-SUZ12 in ESC and sites shared by the proteins in EB (day 8). Occupancy (normalised reads) is indicated by color, according to the scales on the right. At each timepoint, sites are ordered by H3K27me3 in cells expressing SUZ12, from high to low. **b.** H3K27me3 and H4Kac (reads per 10bp window) at the regions shown in Figure 3C in ESC expressing SUZ12, SUZ12Δ93 or JAZF1-SUZ12 at days 0 and 8 after initiation of differentiation into EBs. Dashed box upstream of GATA4 marks the region of ectopic JAZF1-SUZ12 binding from Figure 3C. See also Figure S4.

We also asked whether ectopic JAZF1-SUZ12 binding at non-PRC2 target sites, for example those at *Bax, Cbx3* and *Mcl1*, resulted in ectopic H3K27me3 (Figures 4B, S4A and S4B). However, we found that this was not the case and that H3K27me3 remained low at these sites in cells expressing JAZF1-SUZ12, consistent with the limited amounts of H3K27me3 deposited by this protein at PRC2 target sites.

We next turned our attention to the effects of JAZF1-SUZ12 on H4Kac at PRC2 target sites. We observed an increase in H4Kac at PRC2 binding sites in ESC expressing JAZF1-SUZ12 and this was particularly noticeable at regions of broad SUZ12 occupancy (Figures 4A, 4B and S4A). Cells expressing SUZ12Δ93 also exhibited an increase in H4Kac at these sites, suggesting that much of this effect was due to loss of the repressive function of H3K27me3 rather than due to recruitment of NuA4/TIP60. However, in EBs, JAZF1-SUZ12 caused a marked increase in H4Kac at PRC2 target sites that was not observed for SUZ12Δ93, indicating this was due to fusion to JAZF1 (Figure 4A). Increases in H4Kac in EBs expressing JAZF1-SUZ12 were also observed at individual genes including *T, Pax3, Fgf5, Prdm8* and *Hhex* (Figures 4B and S4A). We did not observe an increase in H4Kac at JAZF1-SUZ12 binding sites shared with JAZF1 (Figure S4B), likely because of endogenous NuA4/TIP60 activity at these sites. We conclude that, in addition to reducing H3K27me3, JAZF1-SUZ12 also increased the active histone mark H4Kac at PRC2 target genes, especially in differentiated cells.

### JAZF1-SUZ12 disrupts gene expression during cell differentiation

We considered that changes in the pattern of PRC2 recruitment and histone modification induced by JAZF1-SUZ12 may be associated with alterations in the pattern of gene expression during ESC differentiation. To address this, we harvested RNA from *Suz12*^GT/GT^ ESC stably expressing GFP, SUZ12, SUZ12Δ93 or JAZF1-SUZ12 before and 4 and 8 days after induction of EB formation. We first determined whether JAZF1-SUZ12 had any effect on the expression of pluripotency genes. We found that *Oct4*, *Fgf4*, *Nanog* and *Utf1* were repressed during EB formation to a similar extent in all of the cell lines, although for *Oct4* and *Fgf4*, repression was delayed in cells containing SUZ12Δ93 or JAZF1-SUZ12 (Figure 5A). These data indicate that the fusion of JAZF1 to SUZ12 has minor effects on silencing of the ESC gene expression program and that this effect is due to loss of the SUZ12 N-terminus.

**Figure 5.**
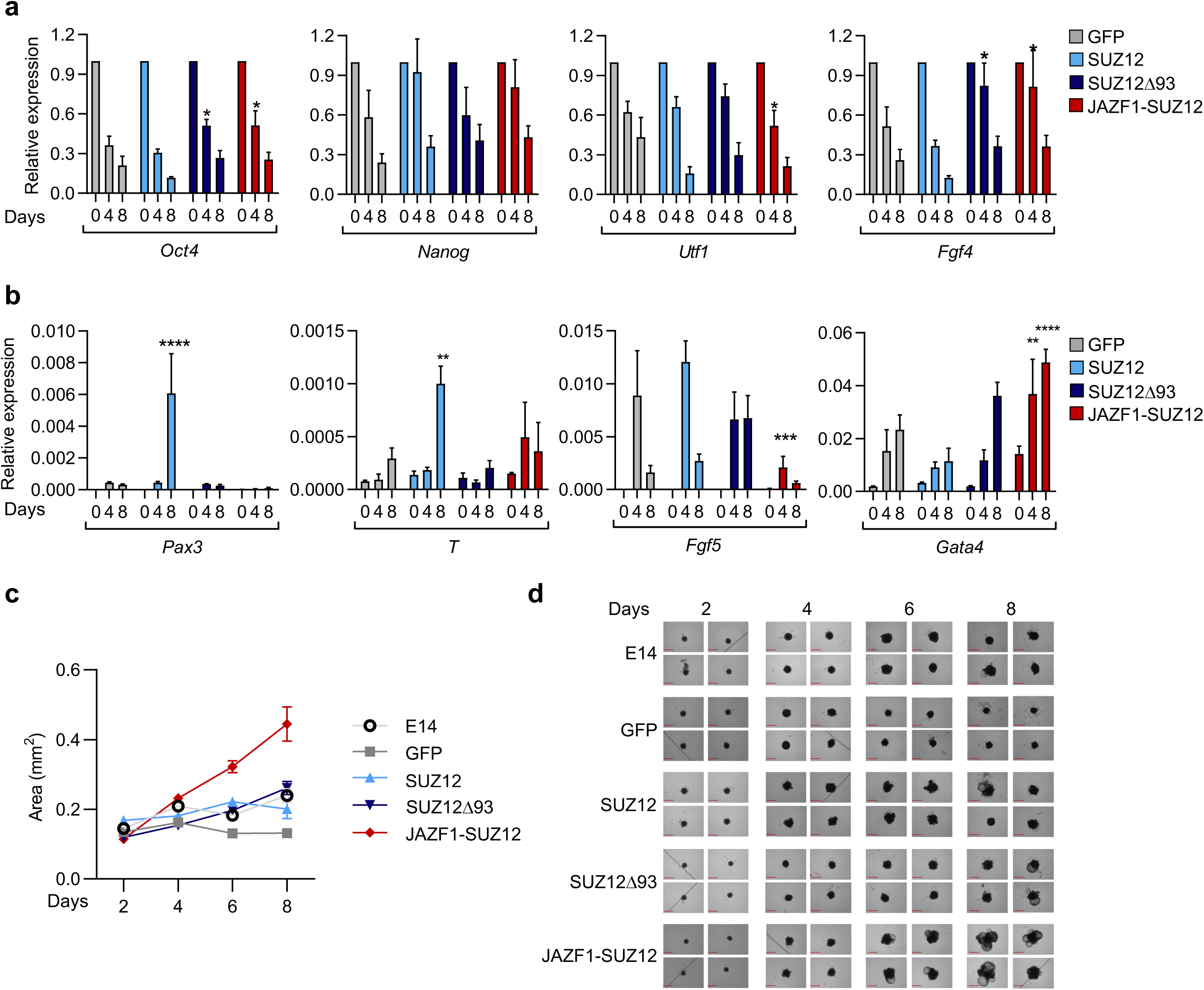
JAZF1-SUZ12 disrupts gene expression during cell differentiation. **a.** Expression of *Oct4*, *Nanog*, *Utf1* and *Fgf4* (relative to *Gapdh* and day 0) at 0, 4 and 8 days after initiation of EB formation in *Suz12*^GT/GT^ ESC expressing GFP, SUZ12, SUZ12Δ93 or JAZF1-SUZ12 (mean and s.d. of 4 independent experiments; *p<0.05, 2-way ANOVA). **b.** Expression of *Pax3, T, Fgf5* and *Gata4* (relative to *Gapdh*) at 0, 4 and 8 days after initiation of EB formation in *Suz12*^GT/GT^ ESC expressing GFP, SUZ12, SUZ12Δ93 or JAZF1-SUZ12 (mean and s.d. of 4 independent experiments; *p<0.05, 2-way ANOVA). **c.** Size of EBs formed by WT E14 ESC and *Suz12*^GT/GT^ ESC expressing GFP, SUZ12, SUZ12Δ93 or JAZF1-SUZ12 measured at 2, 4, 6 and 8 days after initiation of EB formation (mean and s.d, n≥55). **d.** Representative images of *Suz12*^GT/GT^ ESC expressing GFP, SUZ12, SUZ12Δ93 or JAZF1-SUZ12 at 2, 4, 6 and 8 days after initiation of EB formation. The red reference bars mark 0.65 mm.

Given that we had observed dysregulation of SUZ12 recruitment and histone modification during ESC differentiation, we next assessed the effect of JAZF1-SUZ12 on gene induction during EB formation (Figure 5B). We found that JAZF1-SUZ12 disrupted normal patterns of gene induction, but in different ways for different genes. Unlike cells expressing WT SUZ12, cells expressing JAZF1-SUZ12 did not induce *Pax3* during differentiation and also exhibited reduced induction of *Fgf5* and *T*. In contrast, *Gata4* was induced more strongly during differentiation of cells expressing JAZF1-SUZ12. SUZ12Δ93 had similar effects to JAZF1-SUZ12 on *T* and *Gata4* induction, indicating these changes were caused by loss of the SUZ12 N-terminus, whereas the effects of SUZ12Δ93 on *Pax3* and *Fgf5* were distinct to those of JAZF1-SUZ12, indicating these changes were caused by fusion to JAZF1.

We next considered whether these alterations in gene induction in cells expressing JAZF1-SUZ12 were associated with changes in EB formation. To address this, we induced differentiation of WT ESC and *Suz12*^GT/GT^ ESC expressing GFP, SUZ12, SUZ12Δ93 or JAZF1-SUZ12 and measured the size of the resulting EBs after 2, 4 and 8 days (Figures 5C and D). We found that all cell lines formed EBs and, except for cells expressing GFP, these EBs increased in size between days 2 and 8, indicating some rescue of PRC2 function. Strikingly, EBs formed by cells expressing JAZF1-SUZ12 were significantly larger than EBs formed by WT ESC or *Suz12*^GT/GT^ ESC expressing the other constructs. Furthermore, the increase in the size of EBs formed from cells expressing JAZF1-SUZ12 was associated with the formation of multiple cystic cavities (Figure 5D). Production of cystic cavities is associated with expression of endoderm markers such as *Gata4* (Fujikura et al., 2002; Kulinski et al., 2015), consistent with enhanced expression of this gene in EBs containing JAZF1-SUZ12. We conclude that JAZF1-SUZ12 dysregulates gene expression and that this disrupts normal patterns of cell differentiation.

### JAZF1-SUZ12 alters gene expression in primary human endometrial stromal cells

Given that JAZF1-SUZ12 altered gene expression in ESC, we asked whether this effect was also evident in the cell type in which the fusion event occurs; human endometrial stromal cells (hEnSC). To assess this, we transduced primary hEnSC with retroviral vectors encoding SUZ12, JAZF1-SUZ12 or JAZF1, selected for cells expressing the transgenes, and purified RNA before and 4 and 8 days after induction of decidualisation with cyclic AMP (cAMP) and methyl-progesterone acetate (MPA) (Gellersen and Brosens, 2014) (Figures S5A and 5B). By day 8, each hEnSC culture changed from a fibroblast to an epithelioid phenotype and induced expression of the decidualisation marker *PRL* (Figures 6A and S5C). However, at day 0, we found that hEnSC expressing JAZF1-SUZ12 already exhibited increased expression of *IGFBP2* and *ROR2* (Figure 6A), genes that have previously been linked to uterine sarcomas (Cuppens et al., 2015; Przybyl et al., 2018). Thus, JAZF1-SUZ12 disrupts gene expression in primary hEnSC as well as in ESC.

**Figure 6.**
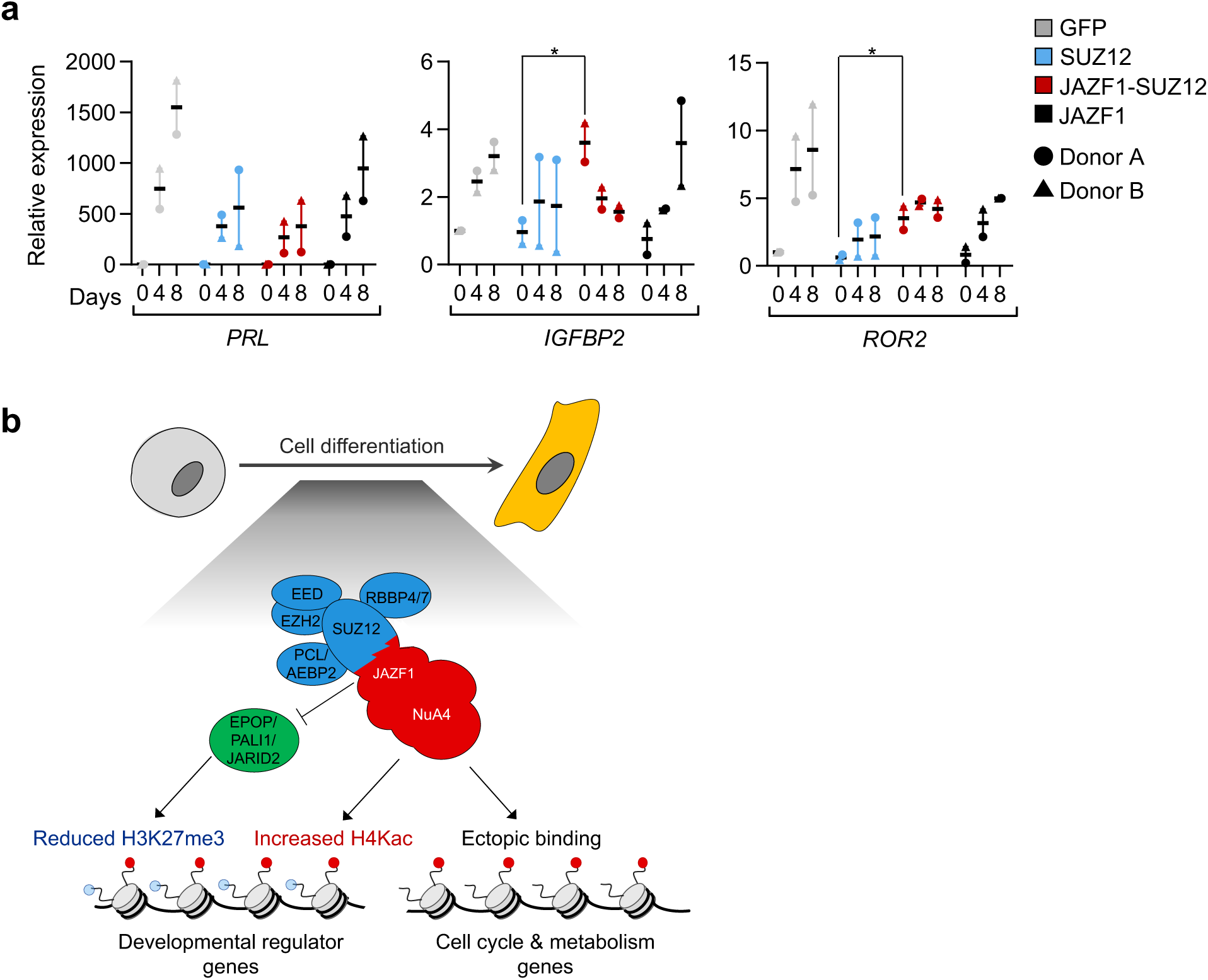
JAZF1-SUZ12 alters gene expression in primary human endometrial stromal cells (hEnSC). **a.** Expression of *PRL, IGFBP2* and *ROR2* (relative to 5S rRNA and GFP day 0) at 0, 4 and 8 days after initiating decidualisation of hEnSC expressing GFP, SUZ12, JAZF1-SUZ12 or JAZF1 (mean and s.d., 2 donors, *p<0.05, t-test). **b.** Model: Loss of the SUZ12 N-terminus prevents interaction of JAZF1-SUZ12 with EPOP, PALI1 and JARID2, while fusion to JAZF1 induces association with NuA4/TIP60. Loss of interaction with EPOP, PALI1 and JARID2 decreases PRC2 occupancy at its target genes and reduces H3K27me3, while fusion to JAZF1 increases H4Kac at these sites during cell differentiation. Fusion to JAZF1 also triggers ectopic SUZ12 recruitment to functional classes of gene with which it does not normally associate. These changes in chromatin occupancy and histone modification are accompanied by dysregulated gene expression and defects in cell differentiation. See also Figure S5.

## DISCUSSION

We have determined the effect of JAZF1-SUZ12 on PRC2 composition, recruitment, histone modification, gene expression and cell differentiation and identified the effects caused by loss of the SUZ12 N-terminus versus those caused by fusion to JAZF1. JAZF1-SUZ12 does not interact with the PRC2.1 subunits EPOP and PALI1 or the PRC2.2 subunit JARID2 and this prevents recruitment from RNA to chromatin. We also confirm that JAZF1-SUZ12 interacts with the NuA4/TIP60 component TRRAP. These alterations in protein interactions are reflected in changes in JAZF1-SUZ12 chromatin occupancy during cell differentiation. In undifferentiated ESC, the pattern of JAZF1-SUZ12 chromatin occupancy resembles WT SUZ12 but becomes more similar to JAZF1 as ESC differentiate into EBs. JAZF1-SUZ12 reduces H3K27me3 and increases H4Kac at PRC2 target genes, and this is coupled with changes in gene expression and morphology during ESC differentiation. JAZF1-SUZ12 also alters gene expression in primary hEnSC. Thus, we conclude that JAZF1-SUZ12 exhibits both loss and gain of function, which leads to dysregulation of gene expression and alterations in cell differentiation (Figure 6B). We propose that these effects of JAZF1-SUZ12 contribute to oncogenesis in cases of LG-ESS with the t(7;17)(p15:q21) rearrangement.

Our results confirm previous observations that JAZF1-SUZ12 does not interact with JARID2 or EPOP (Chen *et al*., 2018) and reveal that this is due to loss of the first 93 amino acids of SUZ12 rather than gain of the JAZF1 N-terminus. The requirement for the first 93 amino acids of SUZ12 for interaction with these proteins is reflected in cryo EM-based reconstructions of PRC2 structure that show that the SUZ12 ZnB (amino acids 76-110) and Zn (amino acids 420-500) regions come together to form a neck-like structure that interacts with JARID2 and is also consistent with the competition observed between JARID2 and EPOP for SUZ12 binding (Chen et al., 2018). We also show that deletion of the N-terminal 93 amino acids abrogates the interaction of SUZ12 with PALI1. That SUZ12 interacts with PALI1 and EPOP through the same region is consistent with the mutual exclusive association of PRC2 with these proteins (Alekseyenko et al., 2014; Hauri et al., 2016). Similarly, that SUZ12 interacts with PALI1 and JARID2 through the same region is consistent with recent findings that PALI1 mimics the allosteric activation function of JARID2 (Zhang et al., 2020).

We found that JAZF1-SUZ12 and SUZ12Δ93 are not recruited to chromatin upon RNA degradation and that this defect is recapitulated in cells lacking JARID2 or EPOP. This suggests that these factors are necessary for the recruitment of PRC2 from RNA to chromatin that occurs upon gene repression (Beltran et al., 2019). We have previously shown that JARID2 is not required for PRC2 RNA binding in cells (Beltran et al., 2016), suggesting that the lack of chromatin recruitment upon RNA degradation reflects a failure in transfer from RNA to chromatin rather than an initial lack of RNA binding. It is interesting that PCL2 is not required for PRC2 recruitment to chromatin upon RNA degradation, even though loss of PCL2 hugely diminishes steady state PRC2 chromatin association (Healy et al., 2019; Højfeldt et al., 2019; Li et al., 2017; Perino et al., 2018). These results suggest that the interaction of PRC2 with chromatin that occurs upon release from RNA is mediated through a different mechanism that relies on JARID2 and EPOP.

Our data reveal how fusion to JAZF alters SUZ12 chromatin occupancy. We found that JAZF1-SUZ12 exhibited a similar binding profile to WT SUZ12 in ESC but as cells differentiate, it takes on a binding profile more similar to that of JAZF1. Thus, fusion of SUZ12 to JAZF1 has a bigger impact on genome localisation in differentiated cells than in undifferentiated cells. This switch in binding preference was not observed for SUZ12Δ93, demonstrating that it depends on fusion to JAZF1 rather than loss of interaction with PRC2 accessory factors. These data are consistent with a model in which cell differentiation is accompanied by changes in chromatin state or changes in the expression of PRC2 and/or NuA4 subunits that shifts the balance of JAZF1-SUZ12 from favouring PRC2 target sites to favouring JAZF1 target sites. We also found that JAZF1-SUZ12 occupied a set of sites that were not bound by SUZ12 or JAZF1 and the increase in the number of these ectopic binding sites during ESC differentiation reflected retention of JAZF1-SUZ12 at sites from which SUZ12 was normally lost.

SUZ12Δ93 also displayed differences in binding and H3K27me3 activity compared with WT SUZ12. The effect of JARID2 (Healy et al., 2019; Højfeldt et al., 2019; Landeira et al., 2010; Oksuz et al., 2018), EPOP (Beringer et al., 2016) or PALI1 (Conway et al., 2018) deficiency on PRC2 genome occupancy or H3K27me3 has been measured previously but the effect of simultaneous loss of all of these interactions provides a comprehensive view of the role of the SUZ12 ZnB/Zn region in PRC2 recruitment and activity. Although JAZF1-SUZ12 and SUZ12Δ93 binding was mostly maintained at broad regions of PRC2 occupancy such as those found at the *Hox* loci, the proteins were reduced at narrower regions of PRC2 occupancy demonstrating a particular requirement for the ZnB/Zn region for PRC2 recruitment to these sites.

Measurement of H3K27me3 and H4Kac at PRC2 target sites revealed that JAZF1-SUZ12-containing PRC2 is less catalytically active than WT SUZ12-containing PRC2; sites with comparable levels of JAZF1-SUZ12 and SUZ12 occupancy exhibit reduced H3K27me3 in cells expressing JAZF1-SUZ12. This was also the case for SUZ12Δ93-containing PRC2, indicating that the reduction of PRC2 catalytic activity in the presence of JAZF1-SUZ12 is due to loss of interaction with JARID2, EPOP and PALI1. This is consistent with the role of JARID2 (Sanulli et al., 2015) and PALI1 (Zhang *et al*., 2020) in the allosteric activation of EZH2. Due to this loss of catalytic activity, ectopic JAZF1-SUZ12 binding was not generally accompanied by ectopic regions of H3K27me3.

Coupled with this decrease in H3K27me3, JAZF1-SUZ12 also increased H4Kac at PRC2 target sites. Increases in H4Kac at PRC2 target genes could also be observed in ESC expressing SUZ12Δ93, indicating that they were primarily due to loss of repressive H3K27me3. However, in EBs, JAZF1-SUZ12 had a more marked effect than SUZ12Δ93 on H4Kac, indicating that this reflected gain-of-function caused by recruitment of NuA4/TIP60 to these sites. Further work will be necessary to determine whether fusion of JAZF1 to SUZ12 also increases other NuA4-mediated chromatin modifications (H2A.Z incorporation, H2A/H2A.Z acetylation) at PRC2 target genes.

The changes in SUZ12 recruitment and histone modification caused by fusion to JAZF1 was reflected in dysregulated patterns of gene expression and differentiation of cells expressing JAZF1-SUZ12. LG-ESS has previously been shown to exhibit elevated expression of polycomb target genes compared with leiomyosarcoma (Przybyl et al., 2018). Thus, our finding that JAZF1-SUZ12 reduces H3K27me3 and increases H4Kac at PRC2 target genes provides an explanation for this elevated expression. Our results further suggest that these changes in H3K27me3 and H4Kac disrupt normal programs of cell differentiation and that this contributes to the role of JAZF1-SUZ12 in oncogenesis. However, confirmation of this hypothesis will require the development of a cell or animal model in which JAZF1-SUZ12 promotes endometrial stromal cell tumourigenesis.

In summary, we have defined the changes in PRC2 composition, recruitment and activity caused by the JAZF1-SUZ12 fusion protein and the effects of these alterations on gene expression and cell differentiation. This work provides new insights into the regulation of PRC2 function and the role of JAZF1-SUZ12 in oncogenesis.

## AUTHOR CONTRIBUTIONS

MT and RGJ designed experiments. MT and MB performed experiments. MT, GK and RGJ analysed data. KV and JM generated reagents. JB and RGJ supervised the research. MT and RGJ wrote the paper.

## ACKNOWLEDGMENTS

We thank the UCL Cancer Institute Genomics Core Facility and UCL Genomics for sequencing. We would like to thank Diego Pasini (Istituto Europeo di Oncologia), Luciano Di Croce (Centre for Genomic Regulation, Barcelona), Adrian Bracken (Trinity College Dublin), Neil Brockdorff (University of Oxford), Amanda Fisher (London Institute for Medical Sciences), Paola Scaffidi (Francis Crick Institute) and Bart Vanhaesebroeck (UCL Cancer Institute) for cell lines and plasmids, and staff in the Implantation Research Clinic at University Hospitals Coventry and Warwickshire NHS Trust for endometrial sample collection. The research was funded by grants from the European Research Council (ERC, 311704) and Worldwide Cancer Research (13-0256) to RGJ, CoNaCyT (411064) to MT and a Wellcome Trust Investigator Award (212233/Z/18/Z) to JJB. GK is a member of the Bill Lyons Informatics Centre and was funded by the Cancer Research UK UCL Centre (award C416/A18088).

## METHODS

### Cell culture

ESC lines were maintained on 0.1% gelatin coated dishes. E14, *Suz12*^GT/GT^ (gift from Diego Pasini), *Ezh*^fl/fl^*, Aebp2*^WT/WT^, *Aebp2*^GT/GT^ (gifts from Neil Brockdorff) and *Jarid2*^GT/GT^ (gift from Amanda Fisher) ESC were cultured in KO-DMEM (ThermoFisher, 10829018), 10% FBS validated for mouse ESC culture (ThermoFisher, 16141079), 5% knockout serum replacement (ThermoFisher, 10828028), non-essential amino acids (ThermoFisher, 11140035), L-glutamine 2 mM (ThermoFisher, 25030-024), 50 μM 2-mercaptoethanol (ThermoFisher, 31350010), 100 U/ml penicillin-streptomycin (ThermoFisher, 15140-122), 1 mM sodium pyruvate (ThermoFisher, 11360039) and 1000 U/ml leukemia inhibitory factor (Stemgent, 03-0011-100). *Pali1*^WT/WT^, *Pali1*^GT/GT^ and *Pcl2*^GT/GT^ *Pcl1/3*^cKO^ (gifts from Adrian Bracken) ESC were maintained in GMEM (Sigma, G5154) with the same supplements, except with no serum replacement and replacing L-glutamine with GlutaMAX (Thermofisher, 35050038), as were *Epop*^GT/GT^ and *Epop*^WT/WT^ cell lines (gifts from Luciano Di Croce) (Beringer et al., 2016) except for 20% FBS. Deletion of *Pcl1* and *Pcl3* was induced by treating *Pcl2*^GT/GT^ *Pcl1/3*^cKO^ cells with 0.5 μM 4-hydroxytamoxifen (Sigma, H6278) for 72 hrs (Healy et al., 2019). NIH-3T3 cells (gift from Bart Vanhaesebroeck) and Lenti-X 293T (Takara Bio Europe) were cultured in high glucose DMEM (Thermofisher, 31966-047), 10% FBS (Thermofisher, 10270-106) and penicillin-streptomycin. Immortalized primary human fibroblasts (gift from Paola Scaffidi) expressing human telomere reverse transcriptase (hTERT) (Scaffidi and Misteli, 2011) were grown in MEM (Thermofisher, 11095-080) supplemented with 15% FBS, 100 U/mL penicillin streptomycin and 2 mM L-glutamine. Drosophila S2 cells (gift from Ivana Bjedov) were grown in Schneider’s Drosophila Medium (ThermoFisher, Cat.no. 21720-02) supplemented with 10% heat-inactivated FBS (ThermoFisher, 10500064) and 25 U/mL of penicillin-streptomycin (ThermoFisher 15140-122). All cell lines were tested negative for mycoplasma (Lonza, LT07-701).

### Embryoid body formation

ESC were differentiated into EBs as described previously (Brien et al., 2012). ESC were filtered using a 70 μm cell strainer, washed twice with LIF-free media and seeded as a single cell suspension on ultra-low attachment T75 flasks (Corning, 3814) at 2×10^6^ cells per flask. LIF-free media was changed every second day and EBs were harvested at days 4 and 8. For quantification of EB growth, 200 cells per well were seeded in 200 μl of LIF-free media in a 96-well ultra-low attachment plate (Costar, 7007) and media was changed every third day. Pictures of single wells were taken with a EVOS FL Auto Imaging System at days 2, 4, 6 and 8, and EB diameter was measured by ImageJ using a macro that used the 650 μm bar in each image as reference.

### Cloning

Human SUZ12 and SUZ12Δ93 were PCR amplified from cDNA. JAZF1 was PCR amplified from IMAGE clone 4814463. To generate JAZF1-SUZ12, nucleotides encoding amino acids 1-128 of JAZF1 and amino acids 94-739 of SUZ12 were PCR amplified and joined in-frame using synonymous *AccIII* sites incorporated into the PCR primers. The constructs were cloned into pCBA-HA and also into pCAG-GFP-2xFLAG (gift from Amanda Fisher). For tagging with Strep-tags, ORFs were PCR amplified from the pCAG plasmids (primers: 5’-TACTTCCAATCCATG and 5’-TATCCACCTTTACTG), C overhangs added with T4 DNA polymerase (M4211), and hybridised with *BaeI*-linearised pCAG-FS2-LIC to which G overhangs had been added. To generate retroviral vectors, ORFs were excised from pCAG-2xFLAG with 5’*EcoRI* and 3’*NotI* and ligated into pMY-IRES-bls.

| Plasmid | Source |
| --- | --- |
| pCAG-GFP(FLAG) | A. Fisher |
| pCAG-SUZ12(FLAG) | This work |
| pCAG-SUZ12D93(FLAG) | This work |
| pCAG-JAZF1-SUZ12(FLAG) | This work |
| pCAG-JAZF1(FLAG) | This work |
| pCAG-(FS2)GFP | This work |
| pCAG-(FS2)SUZ12 | This work |
| pCAG-(FS2)SUZ12D93 | This work |
| pCAG-(FS2)JAZF1-SUZ12 | This work |
| pCAG-(FS2)JAZF1 | This work |
| pMY-IRES-bls-GFP(FLAG) | This work |
| pMY-IRES-bls-SUZ12(FLAG) | This work |
| pMY-IRES-bls-JAZF1-SUZ12(FLAG) | This work |
| pMY-IRES-bls-JAZF1(FLAG) | This work |
| pCBA-SUZ12 (HA) | This work |
| pCBA-SUZ12D93 (HA) | This work |
| pCBA-JAZF1-SUZ12 (HA) | This work |
| pCBA-JAZF1 (HA) | This work |
| pCAG-(FS2)AEBP2 | N. Brockdorff; Grijzenhout et al., 2016 |
| pCAG-(FS2)EPOP | N. Brockdorff; Grijzenhout et al., 2016 |
| pLenti-(FLAG/HA) PALI1 | A. Bracken; Healy et al., 2019 |
| PCL3-V5 | J. Reiter; Hunkapiller et al., 2012 |

### Transfection and generation of cell lines

pCAG constructs were transfected into *Suz12*^GT/GT^ ESC with Effectene (Qiagene, 301425), following the manufacturer’s protocol, cells selected with 2 μg/ml puromycin and stable cell lines expanded from single cell colonies. NIH3T3 cells were transfected with 1 μg/ml plasmid with polyethyleneimine (3 μg/ml; Polysciences, 23966). Media was changed the next day and cells harvested 48 hrs later. Retroviruses were produced by transfection of lenti-X 293T cells with 1.5 μg of pMY-IRES-bls constructs, 1 μg of pCMVi and 1 μg of pMDG (gifts from Greg Towers) using Fugene-HD (Promega, E2311). Virus was collected from 4x 10 cm plates 48, 72 and 96 hrs after transfection and concentrated at 17,000 x g for 2 hrs at 4°C. The viral pellet was resuspended in 1 ml FBS-free DMEM/F12 and then supplemented with 10% DCC FBS prior to freezing at −80°C. To make the FLAG-SUZ12 spike-in control material for ChIP-seq, hTERT cells were transduced with 1.5 ml of concentrated pMY-SUZ12-FLAG-IRES-bls virus in the presence of 8 μg/ml polybrene (Merck, TR-1003-G). Cells were then spinoculated at 500 x g for 1 hr at RT and incubated for 12 hrs. Cells were washed with fresh media and, after 3 days, selected with 5 μg/ml blasticidin.

### hEnSC cell culture, transduction and decidualisation

The collection of endometrial biopsies was approved by the NHS National Research Ethics - Hammersmith and Queen Charlotte’s & Chelsea Research Ethics Committee (REC reference: 1997/5065) and Tommy’s National Reproductive Health Biobank (REC reference: 18/WA/0356). Samples were obtained using a Wallach Endocell sampler 5 to 10 days after the pre-ovulatory luteinizing hormone (LH) surge. hEnSC were harvested as previously described (Barros, Brosens and Brighton, 2016). Briefly, endometrial biopsies were subjected to enzymatic digestion using 500 µg/ml collagenase type Ia (Sigma-Aldrich,) and 100 µg/mL DNase I (Lorne Laboratories Ltd) for 1 hr at 37°C. Digested tissue was filtered through a 40 µm cell strainer to remove glandular cell clumps, and the flow-through collected and cultured in DMEM/F12 (Thermo Scientific, 31330038) containing 10% dextran-coated charcoal-treated foetal bovine serum (DCC-FBS), 1 × antibiotic-antimycotic mix (ThermoFisher, 15240062), 10 µM L-glutamine, 1 nM estradiol (Sigma, E2758) and 2 μg/mL insulin (Santa Cruz, sc-360248). Cells were reseeded at a 1:2 or 1:3 ratio when confluence was reached. All experiments were carried out before reaching the 10^th^ passage.

hEnSC were seeded at a confluency of 10^5^ cells/well in 6 well plates and 1.5 ml of concentrated virus added to each well in the presence of 8 μg/ml polybrene (Merck, TR-1003-G). Cells were then spinoculated at 500 x g for 1 hr at RT and incubated for 12 hrs. Cells were washed with fresh media and, after 3 days, selected with 5 μg/ml blasticidin until they reached near confluency and no cells were found in the non-transduced control plate. Cells were reseeded and, at confluency two days later, decidualised for up to 8 days in DCC media with 2% DCC FBS supplemented with antibiotic antimycotic solution, l-glutamine, 50 μM 8-Bromoadenosine 3’, 5’ cyclic mono-phosphate (8-Br-cAMP, Merck B5386) and 1 μM medroxyprogesterone acetate (MPA, Sigma M1629), with the media changed every second day.

### Co-immunoprecipitation and Strep-Tactin pull-down

Immunoprecipitations were performed as described (Conway et al., 2018), with minor modifications. 50 million cells per IP were suspended in 500 µl of high salt buffer (50 mM Tris-HCl, pH 7.2, 300 mM NaCl, 0.5% (v/v) NP-40, 1 mM EDTA pH7.4, Complete protease inhibitor (Roche, 11873580001) and 1 mM DTT) and sonicated for 3x 10 seconds using a Bioruptor Pico (Diagenode). Cells were then rotated at 4°C for 20 mins before the lysates were diluted with 500 µl of no salt buffer (50 mM Tris-HCl, pH 7.2, 0.5% (v/v) NP-40, 1 mM EDTA pH7.4, Complete protease inhibitor and freshly added 1 mM DTT). Afterwards, lysates were clarified by centrifugation for 10 min at 17,000 x g at 4°C and 50 μl taken from the sample for use as input. The remainder of the lysate was incubated with 2 µg of anti-FLAG M2 (Sigma, F1804), SUZ12 (CST, 3737) or anti-V5 (Abcam, ab15828) antibody with 250 U/mL benzonase (Sigma, E1014) for 16 hrs. 50 μl Protein G Dynabeads (Invitrogen, 10003D) per sample were washed 3 times with wash buffer (1:1 dilution of high salt: no salt buffer), resuspended in their initial volume, and incubated with the protein lysates for 2 hrs with rotation at 4°C. The flow-through was removed and immunocomplexed beads washed 5 times with 1 ml of wash buffer. Beads were resuspended in 100 μl of a 1:1 mix of 2X Laemmli buffer (4% SDS, 240 mM Tris pH 6.7, 2% beta-mercaptoethanol, 20% glycerol and 0.2% bromophenol blue) and 10 mM Tris HCl, while 50 μl of 2X Laemmli buffer was added to input samples and both heated at 95°C for 5 min. Finally, 10 μl of each IP and input sample were resolved by SDS-PAGE.

Strep-Tactin pull-down was carried out as for FLAG immunoprecipitation, with the following modifications. To block binding of any unspecific biotinylated proteins, 10 μg/ml of avidin (IBA, Cat No. 2-0204-015) was added to the extracts after cell lysis. Cell extracts were then incubated for 30 min at 4°C with rotation and centrifuged at 20,817 x g for 5 min at 4°C and the supernatant recovered. Supernatant was added to 10 μl of StrepTactin superflow high-capacity resin (IBA, Cat no. 2-1208-002) and incubated with rotation for 4 hrs at 4°C. Resin was then washed 5 times with wash buffer, each time pelleting the resin at 1000 x g for 5 mins at 4°C. Protein was eluted by boiling the resin in Laemmli buffer.

### Nuclear fractionation and RNaseA treatment

RNaseA treatment and cell fractionation was performed as described (Beltran et al., 2016; Zoabi et al., 2014). ESCs were trypsinized, washed twice with PBS, permeabilized with 0.05% Tween-20 in PBS for 10 mins on ice, washed once, resuspended with PBS and mock-treated or treated with 1 mg/ml RNaseA (Sigma, R6513) for 30 mins at RT. Cells were centrifuged at 1200 rpm, washed twice and resuspended in 1 ml of buffer A (10 mM HEPES (pH 7.9), 10 mM KCl, 1.5 mM MgCl2, 0.34 M sucrose, 10% glycerol, 1 mM DTT with Complete protease inhibitor). Triton X-100 (0.1%) was added, and the cells were incubated for 5 mins on ice. Nuclei were collected by low-speed centrifugation (4 mins, 1,300 x g, 4°C). The supernatant (cytoplasmic fraction) was further clarified by high-speed centrifugation (15 min, 20,000 x g, 4°C). Nuclei were washed twice in buffer A, and then lysed in buffer B (3 mM EDTA, 0.2 mM EGTA, 1 mM DTT, Complete protease inhibitor). Insoluble chromatin was collected by centrifugation (4 min, 1,700 x g, 4°C), and the supernatant (nucleoplasmic fraction) was recovered. The final chromatin pellet (chromatin fraction) was washed twice with buffer B, resuspended in 1X Laemmli buffer, sonicated (Diagenode Bioruptor Pico) and resolved by SDS-PAGE.

### Immunoblotting

Proteins were resolved along side PageRuler (ThermoFisher, 26620) by SDS-PAGE using the Mini-PROTREAN Tetra Cell system (BioRad) in 200 mM glycine, 24 mM Tris base and 0.1% SDS. Proteins were transferred to 0.45 μM nitrocellulose membrane (GE Healthcare, 15269794) using a Mini Trans-Blot system (Biorad, 1610158) at 350 mA for 2 hrs. Membranes were blocked with 5% non-fat dried milk plus 0.1% Tween (Sigma, P1379) in TBS (TBST) for 1 hr at RT. Proteins were detected with primary antibodies to FLAG M2 (Sigma, A8592), HA 3F10 (Roche, 12013819001), V5 (Abcam, ab15828), SUZ12 (Santa Cruz sc-46264), EZH2 (CST, 3147), JARID2 (CST, 13594), AEBP2 (CST 14129), PCL2 (Proteintech 16208-1-AP), EPOP (kind gift of L. Di Croce), TRRAP (Abcam, ab73546), FUS (Novus Biologicals 100-565), β-actin (CST 4967), alpha tubulin (CST 2144), H3K27me3 (Abcam ab192985) and H3 (Abcam ab1791) and HRP-conjugated secondary antibodies (anti-mouse (Dako, P0447) or anti-rabbit (Dako, P0448)). Proteins were visualised using the Clarity ECL Western Substrate (Biorad, 1705061) and detected using an ImageQuantLAS 4000 imager and ImageQuantTL software (GE). Contrast and brightness were altered in a linear fashion equally across the whole image.

### RNA quantification

RNA was purified using TRIsure (Bioline, BIO-38033) and reverse transcribed using the Im-Prom-II Reverse Transcription System (Promega, A3800) and random hexamer primers. Specific RNAs were quantified using the QuantiTect SYBR Green PCR Kit (Qiagen, 204145) and a QuantStudio 5 Real-Time PCR System (Thermofisher) with the primers shown below.

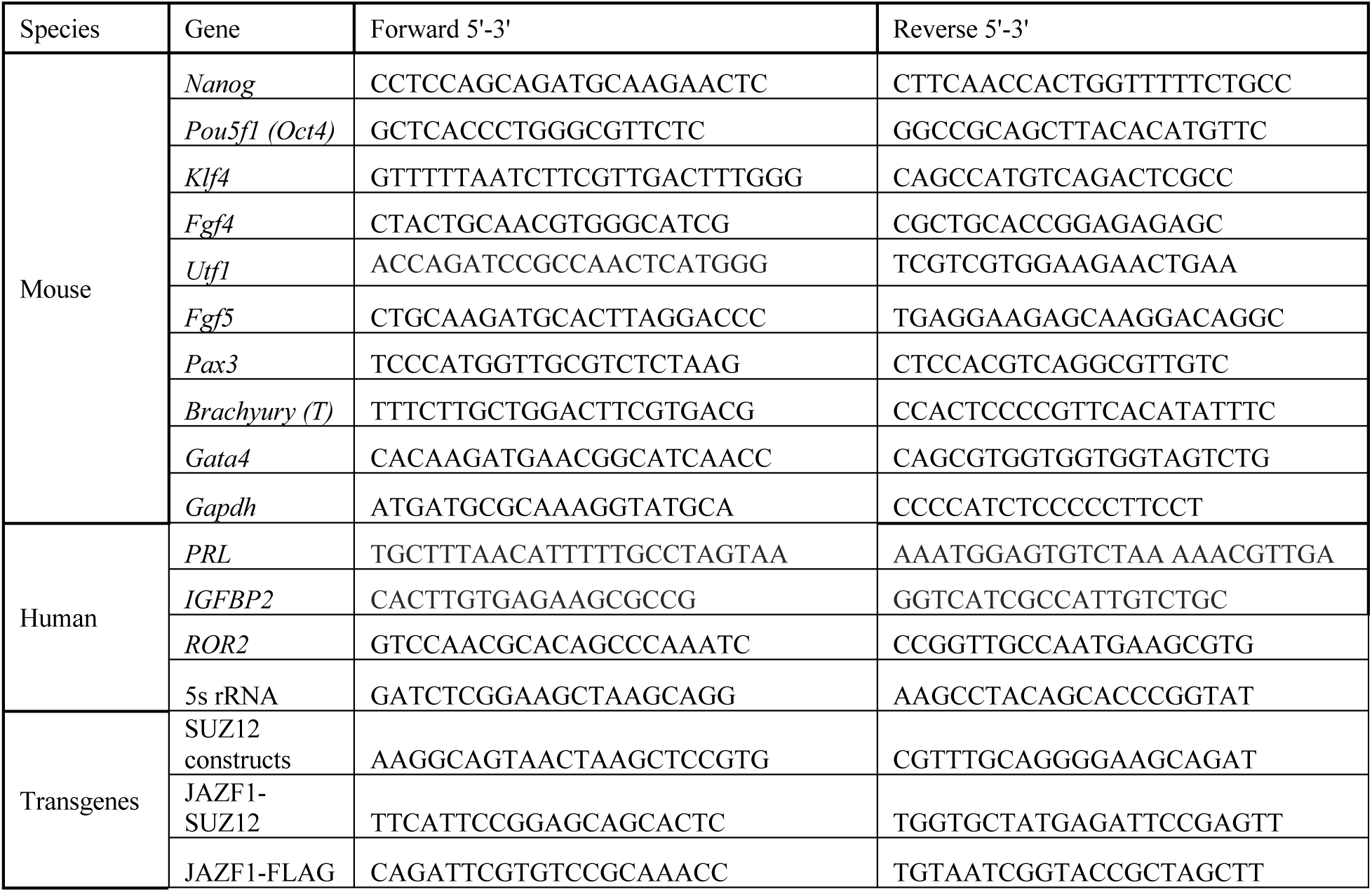

### Calibrated ChIP-sequencing (cChIP-Seq)

All ChIP-seq experiments were performed in duplicate. Cells were trypsinised, resuspended in PBS and then crosslinked by adding 1/10 volume of cross-linking solution (11% formaldehyde, 0.1 M NaCl, 1mM EDTA pH 8, 0.5 mM EGTA pH 8, 50 mM HEPES pH 8) for 15 min. Formaldehyde was quenched with 1.25 mM glycine. Cells were washed twice with ice-cold PBS, centrifuging at 290 x g at 4°C for 10 min each time, flash-frozen and stored at −80°C. Cells were thawed, resuspended in lysis buffer 1 (50 mM HEPES pH 7.5, 140 mM NaCl, 1 mM EDTA, 10 % glycerol, 0.5 % IGEPAL CA-630, 0.25 % Triton X-100) and incubated at 4°C for 10 mins with rocking. Cells were recovered by centrifugation at 290 x g for 10 min at 4°C and nuclei resuspended in the same volume of lysis buffer 2 (10 mM Tris pH 8, 200 mM NaCl, 1 mM EDTA, 0.5 mM EGTA, supplemented with 0.1 DTT and Complete protease inhibitor) and incubated at 4°C for 10 mins with rocking. Nuclei were pelleted again at 290 x g for 10 mins at 4°C and resuspended in 100 μl of lysis buffer 3 (10mM Tris pH 8, 100mM NaCl, 1mM EDTA, 0.5mM EGTA, 0.1% sodium deoxycholate, 0.5% N-lauryl sarcosine, 0.2 % SDS, supplemented with 0.1 DTT and Complete protease inhibitor) per 5×10^6^ cells. Cells were incubated for 30 mins on ice and sonicated for 15 cycles of 30s on and 30s off using a Diagenode Bioruptor Pico. 2×10^7^ ESC or the EBs formed from 2x 10 cm plates (4 day timepoint) or 1x 10 cm plate (8 day timepoint) were used per ChIP. An aliquot of the sonicated chromatin was reverse crosslinked by heating at 65°C for 1 hr, treated with 0.5 mg/ml RNaseA and 0.2 mg/ml of proteinase K (Ambion, AM2546). DNA was purified from this aliquot using KAPA Pure beads (Roche, KK8000), resuspended in 10 mM Tris pH 8 and quantified using the Qubit dsDNA HS Assay kit (Thermofisher, Q32851). Each ChIP was then performed with mouse chromatin equivalent to 45 μg of purified DNA. For ChIPs for FLAG-tagged proteins, sonicated human chromatin (extracted from hTERT cells stably expressing FLAG-SUZ12) equivalent to 5 μg of purified DNA was added as a calibration control. For ChIPs for histone modifications, *Drosophila* chromatin (extracted from S2 cells) equivalent to 5 μg of purified DNA was added. The lysates were topped up to 900 μl per IP, 100 μl Triton X-100 10% solution added, and insoluble material removed by centrifugation at 17,000 x g for 20 mins at 4°C. 2% of the lysate was stored at −20°C to be used as input and the remaining lysate was incubated overnight at 4°C with 50 μl protein G Dynabeads preincubated for at least 4 hr with 2.5 μg of anti-FLAG M2 (Sigma, A8592), H3K27me3 (Abcam, ab192985) or H4 pan-acetyl (Thermofisher, PA5-40083) antibody. Beads were washed six times with wash buffer (50 mM Hepes, 1 mM EDTA, 0.5 M LiCl, 0.7% sodium deoxycholate, 1% NP-40) and once with TE with 50 mM NaCl and bound complexes eluted in elution buffer (50 mM Tris, 10 mM EDTA, 1% SDS). Crosslinks were reversed by heating at 65°C for 8 hrs and DNA purified as before.

Sequencing libraries were generated from 1.2 ng of DNA using the NEBNext Ultra II DNA library preparation kit (NEB, E7645) with 9 cycles of PCR. Library quality and size distribution was assessed using the 2100 Bioanalyzer High Sensitivity DNA Kit (Agilent, 5067-4626) followed by qPCR quantification with the Kapa Library Quant Kit (Roche, KK4824). FLAG libraries were subjected to 75 bp single-end sequencing on an Illumina NextSeq 550 platform and histone modification libraries were subjected to 138 bp single-end sequencing on an Illumina NovaSeq platform.

### ChIP-seq data analysis

GFP, SUZ12, SUZ12Δ93, JAZF1-SUZ12 and JAZF1 ChIP-seq reads were aligned to a concatenated genome sequence of mouse (mm10) and human (hg19) using bowtie2 with the -- very-sensitive option (Langmead and Salzberg, 2012). Uniquely mapped reads were extracted using samtools (Li et al., 2009) and were used for the downstream analysis. For each sample, endogenous (mouse) and exogenous (human) reads were segregated into two bam files using samtools.

Spike-in calibration was performed as previously described (Fursova et al., 2019) and down-sampled bam files were generated for the endogenous mouse data. The two replicates for the endogenous data were merged and peaks were called using MACS2 with the --broad option and - -broad-cutoff = 0.0001 (Feng et al., 2012). Peaks overlapping mouse blacklist regions (Amemiya et al., 2019) or peaks called in any of the GFP control datasets were removed using bedTools (Quinlan and Hall, 2010). Genome coverage tracks were obtained using MACS2 pileup function, which were then converted to bigwigs and visualized with the UCSC Genome Browser (Kent et al., 2002).

A combined peak-set was created from the merged peaks from all the factors at the three time-points using DiffBind in R with the default parameters (Ross-Innes et al., 2012). Correlations between the peaks in different samples were calculated with DiffBind using the Pearson correlation co-efficient. Peak overlaps were calculated with dba.overlap and UpSet plots generated with UpSetR (Conway et al., 2017). Metaplots and heatmaps were generated with computeMatrix and plotProfile/plotHeatmap functions from deepTools (Ramírez et al., 2014).

H3K27me3 and H4Kac ChIP-seq reads were aligned to a concatenated genome sequence of mouse (mm10) and *Drosophila* (BDGP5.25) using bowtie2 with the --very-sensitive option. Uniquely mapped reads were extracted and separated into endogenous (mouse) and exogenous (*Drosophila*) reads using samtools. The normalization factor for the mouse bams was calculated using the *Drosophila* spike-in according as previously described (Orlando et al., 2014). The normalization factor was used to generate the mouse bigwigs using the bamCoverage function from deepTools.

The closest gene transcription start sites (TSS; Ensembl v98) to each SUZ12, JAZF1-SUZ12 and JAZF1 MACS peak was identified with bedTools. Gene TSS within 1 kb of a peak were considered to be bound. Gene Ontology terms enriched in the sets of genes bound by each factor were identified using g:Profiler with the default settings (Raudvere et al., 2019).

### Statistical analysis

Measurements of chromatin association after RNA degradation were performed in triplicate and mean and standard deviation plotted in GraphPad Prism (GraphPad Software, USA) The significance of changes in chromatin association were estimated using a two-tailed Student’s t-test. Measurements of relative gene expression during EB formation were performed in quadruplicate, mean and standard deviation plotted and the significance of differences between samples were estimated using 2-way ANOVA using GraphPad Prism. Mean and standard deviation of EB size was plotted and the significance of differences between samples estimated using 2-way ANOVA (E14 n=95, GFP n=86, SUZ12 n=86, SUZ12Δ93 n=96, JAZF1-SUZ12 n=55. Measurements of relative gene expression in hEnSC were performed in duplicate (cells from two different donors) and the significance of differences between samples were estimated using Student’s t-test.

---

**Figure S1.**
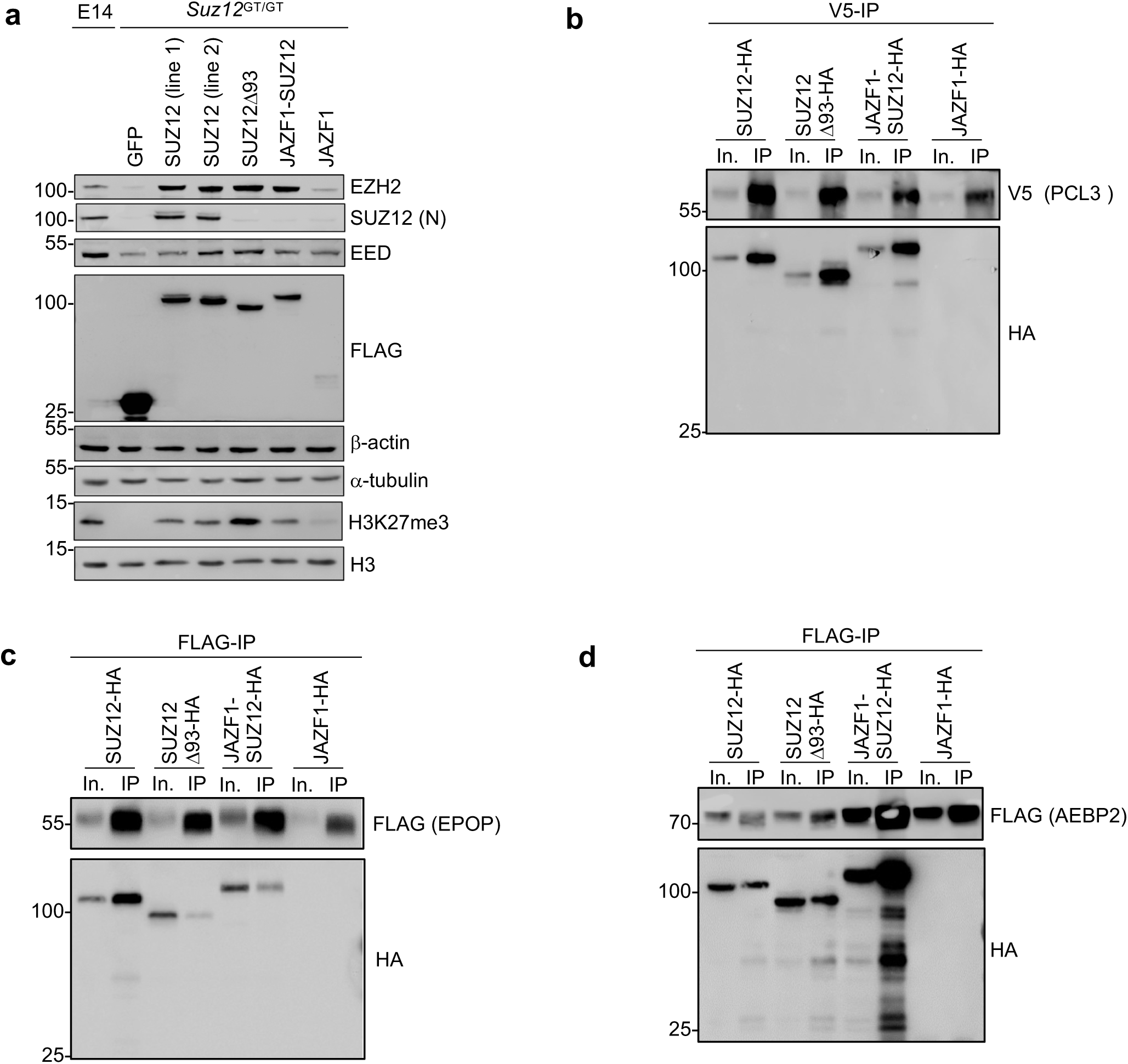
JAZF1-SUZ12 interacts with PCL3 and AEBP2 but shows reduced interaction with EPOP. **a.** Levels of FLAG-tagged GFP, SUZ12 (2 different clones), SUZ12Δ93, JAZF1-SUZ12 and JAZF1 in stable *Suz12*^GT/GT^ cell lines in comparison to WT E14 cells. **b.** Immunoblots for V5 and HA in input (In.) and SUZ12 IP fractions from NIH-3T3 cells co-transfected with V5-tagged PCL3 and HA-tagged SUZ12/JAZF1 constructs. **c.** As b., except in cells transfected with FS2-EPOP and HA-tagged SUZ12/JAZF1 constructs. **d.** As b., except in cells transfected with FS2-AEBP2 and HA-tagged SUZ12/JAZF1 constructs.

**Figure S2.**
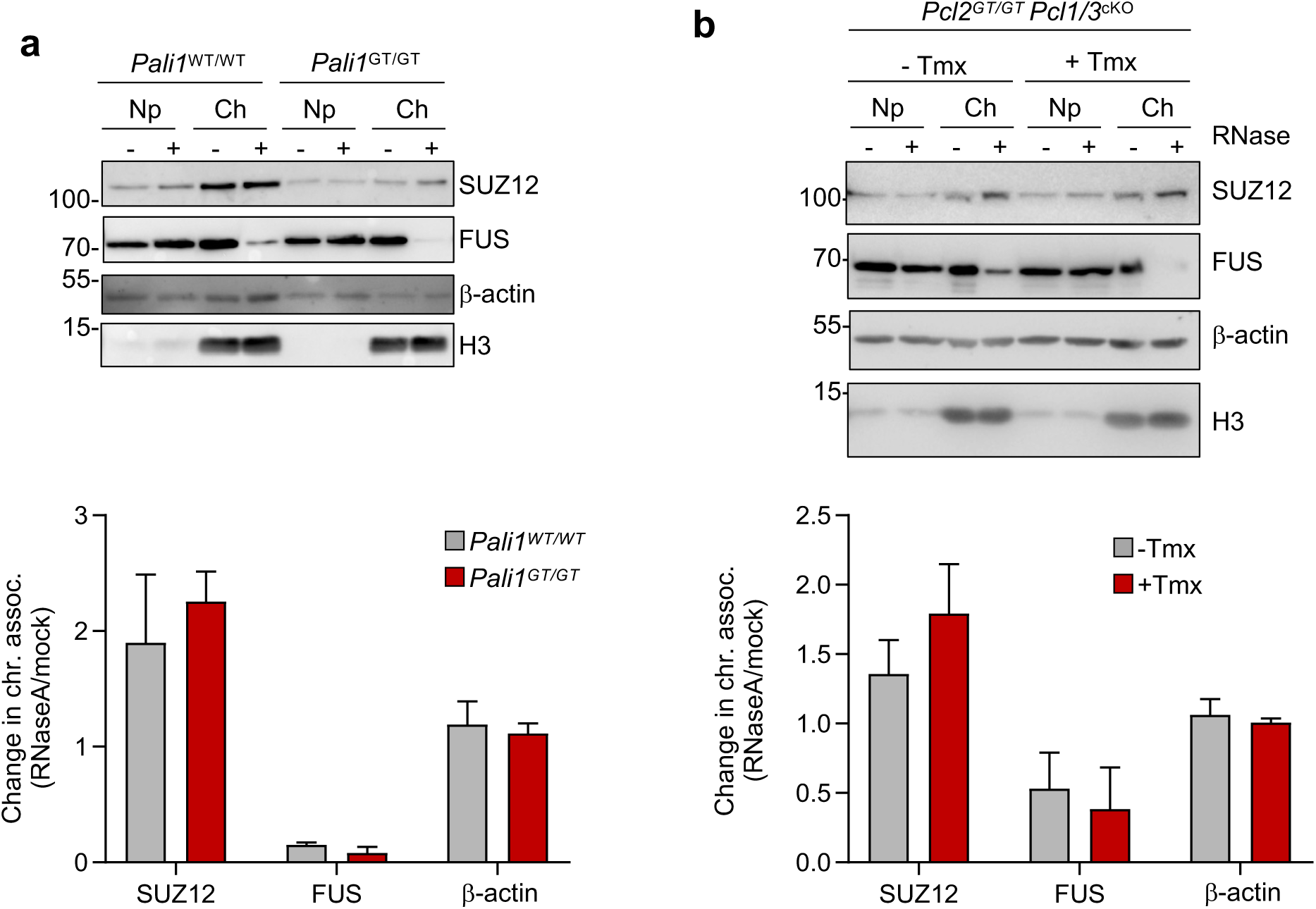
PALI1 and PCL1-3 are not required for recruitment of PRC2 to chromatin upon RNA depletion. **a.** *Left:* Immunoblots for SUZ12, FUS, β-actin, and H3 in nucleoplasm (Np) and chromatin (Ch) fractions from mock or RNaseA-treated WT or *Pali1*^GT/GT^ cells. Representative of two independent experiments. *Right*: Fold change in FLAG, FUS and β-actin in the chromatin fraction upon RNaseA treatment (mean and s.d, n=2) **b.** Immunoblots for SUZ12, FUS, β-actin and H3 in nucleoplasm (Np) and chromatin (Ch) fractions from mock or RNaseA-treated *Pcl2*^GT/GT^ *Pcl1/3*^cKO^ cells pre-treated or not with 4-hydroxytamoxifen (tmx) to induce deletion of *Pcl1* and *Pcl3* (mean and s.d., n=2).

**Figure S3.**
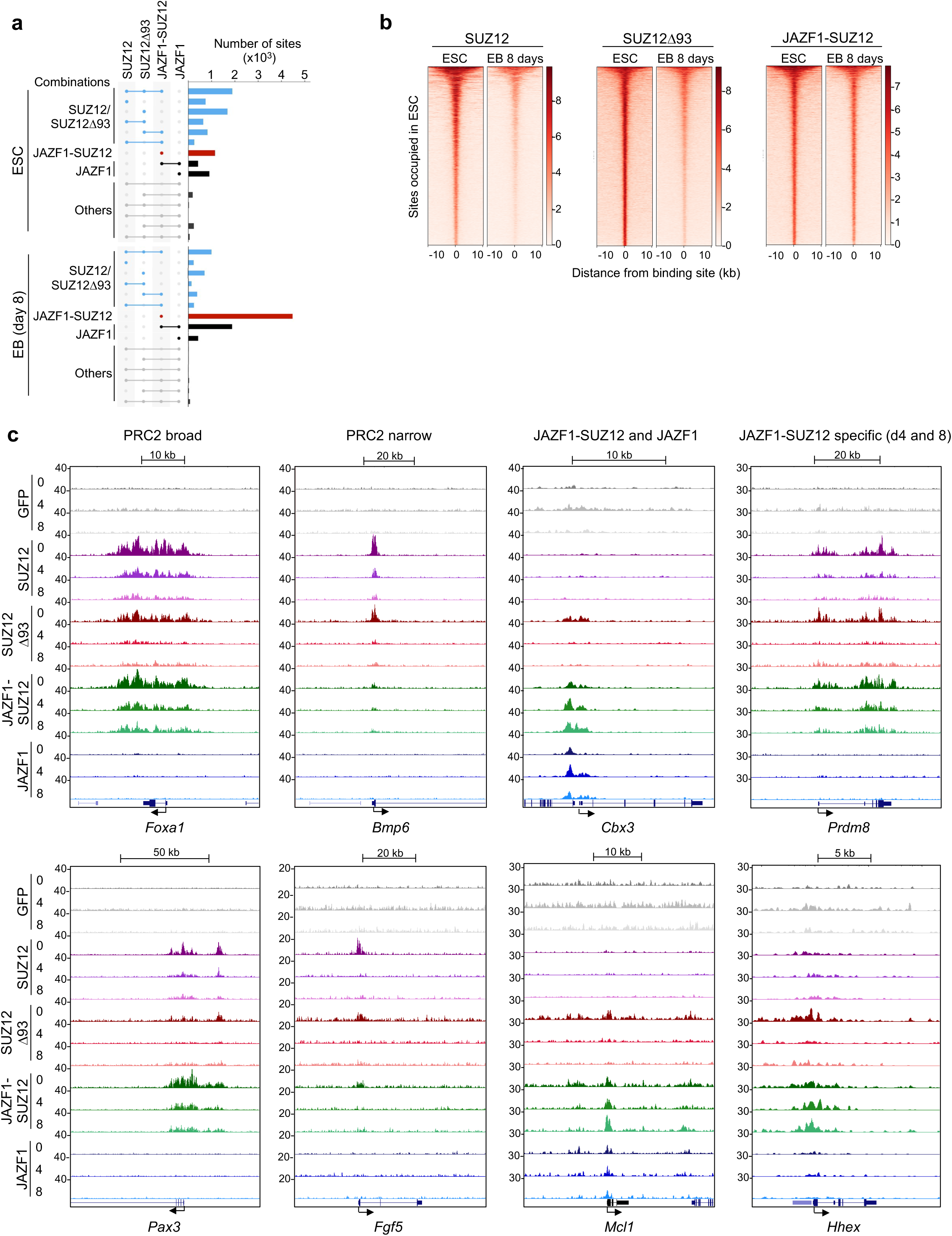
Differences in chromatin occupancy between JAZF1-SUZ12 and SUZ12. **a.** UpSet plots showing the number of sites bound by different combinations of proteins in ESC and in EBs (8 days after induction of ESC differentiation). Binding combinations are grouped into sites bound by SUZ12 and/or SUZ12Δ93 (light blue), sites bound by JAZF1-SUZ12 alone (red), sites bound by JAZF1-SUZ12 and/or JAF1 (black), and other combinations (grey). **b.** Heatmaps showing SUZ12, SUZ12Δ93 or JAZF1-SUZ12 occupancy in ESC and day 8 EBs at sites occupied by each factor in ESC. Occupancy (normalised reads) is indicated by color, according to the scales on the right. For each factor, binding sites are ordered by occupancy in ESC, from high to low. **c.** Further examples of the 4 patterns of JAZF1-SUZ12 occupancy shown in Figure 3C. Left: broad regions of SUZ12 binding at *Foxa1* and *Pax3.* Centre left: narrow regions of SUZ12 binding at *Bmp6* and *Fgf5.* Center right: Shared JAZF1-SUZ12 and JAZF1 occupancy at *Cbx3* and *Mcl1.* Right: JAZF1-SUZ12-specific binding at *Prdm8* and *Hhex* in day 4 and day 8 EBs.

**Figure S4.**
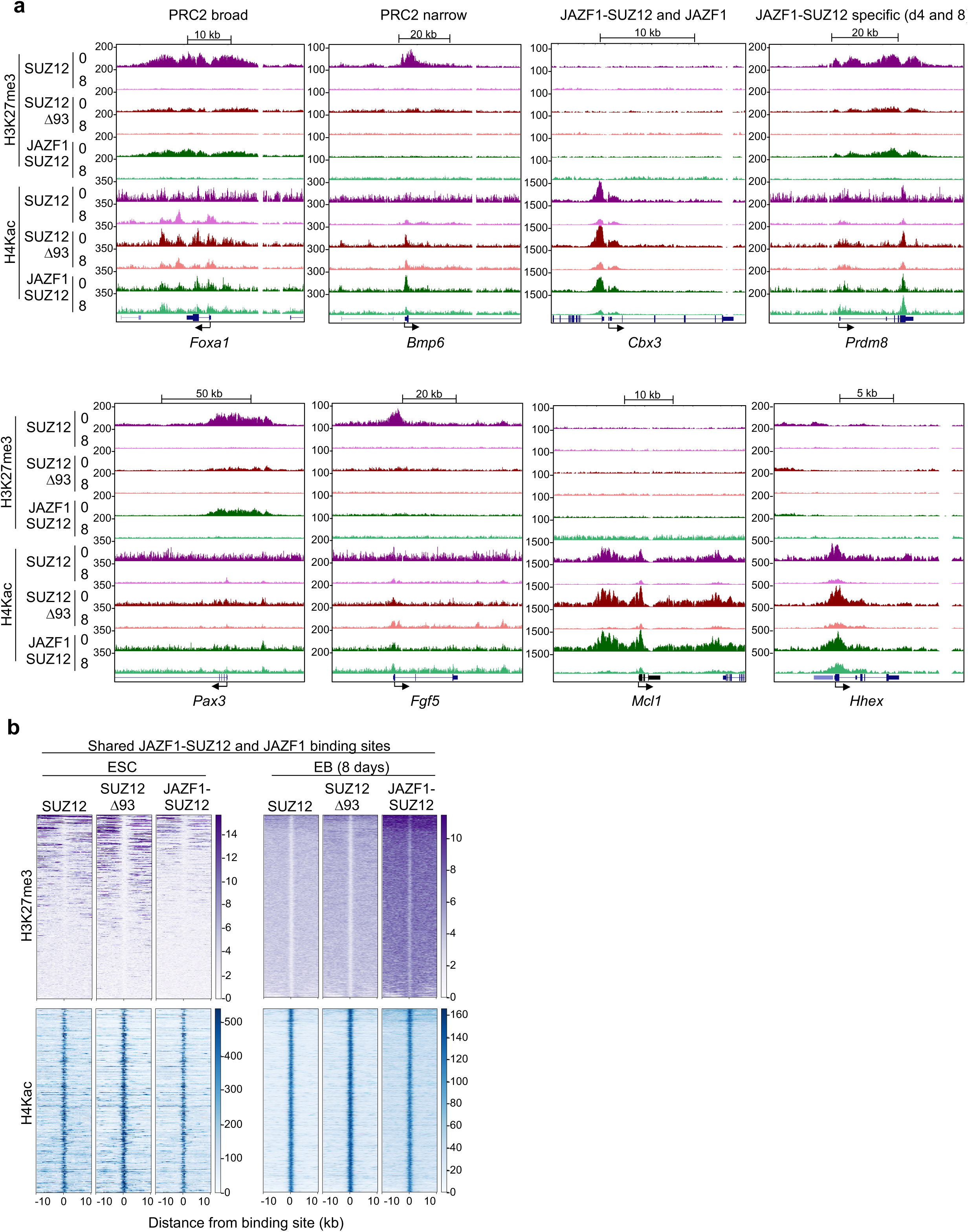
Changes in H3K27me3 and H4Kac in cells expressing JAZF1-SUZ12. **a.** H3K27me3 and H4Kac (reads per 10bp window) at the regions shown in Figure S3C in ESC expressing SUZ12, SUZ12Δ93 or JAZF1-SUZ12 at days 0 and 8 after initiation of differentiation into EBs. **b.** Heatmaps of H3K27me3 (purple, top) and H4Kac (blue, bottom) around binding sites shared by JAZF1 and JAZF1-SUZ12 in ESC and in EB (day 8). Occupancy (normalised reads) is indicated by color, according to the scales on the right. At each timepoint, sites are ordered by H3K27me3 occupancy in cells expressing SUZ12, from high to low.

**Figure S5.**
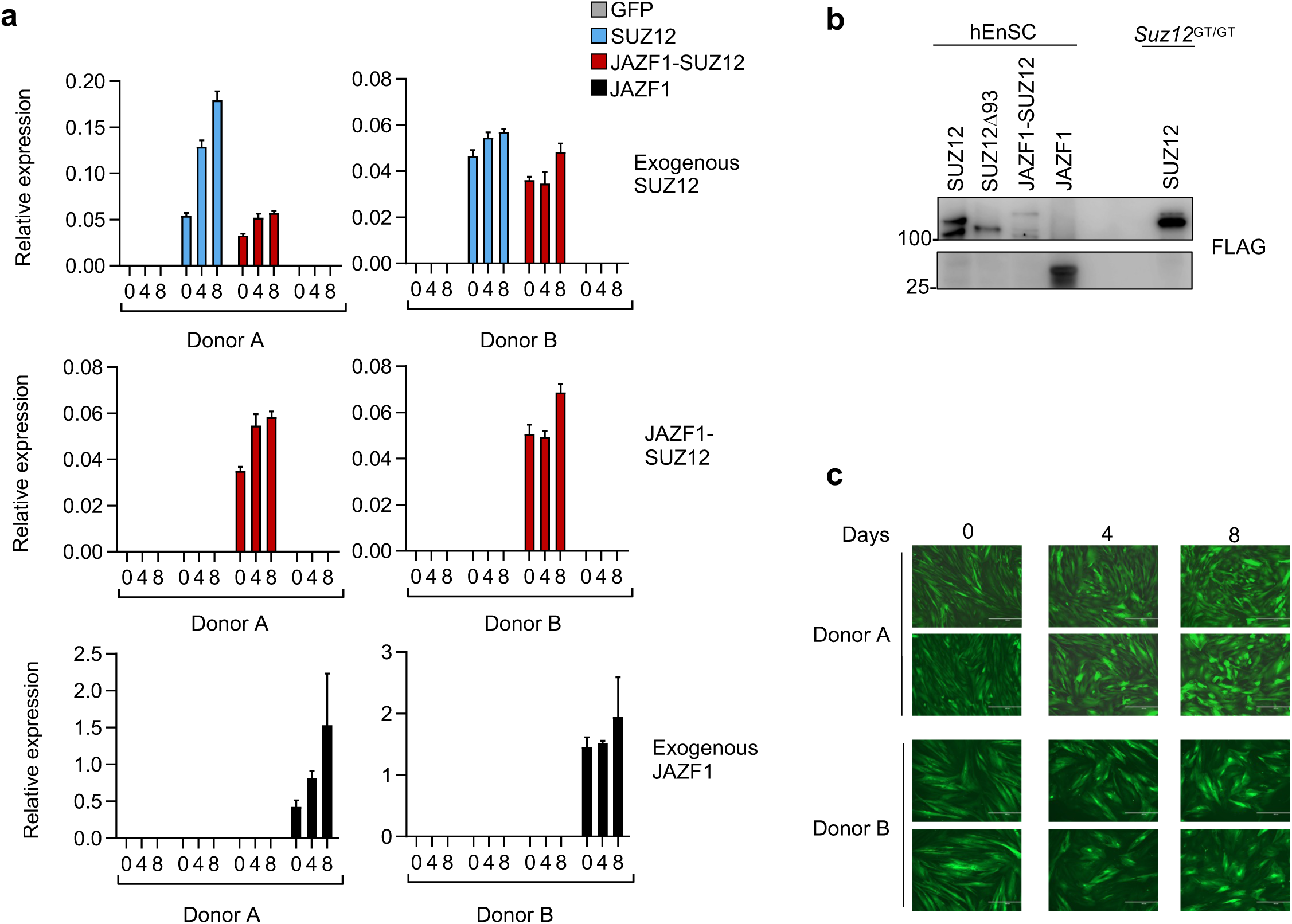
Expression of transgenes in decidualised hEnSC. **a.** Expression of exogenous *SUZ12, JAZF1-SUZ12* and *JAZF1* constructs relative to 5S rRNA at 0, 4 and 8 days after initiating decidualisation of hEnSC from 2 donors. **b.** Immunoblot for FLAG-tagged proteins in whole cell lysate of hEnSC stably expressing FLAG-tagged SUZ12, SUZ12Δ93, JAZF1-SUZ12 and JAZF1 in comparison to *Suz12*^GT/GT^ ESC stably expressing FLAG-tagged SUZ12. **c.** Representative images of hEnSC from two donors expressing GFP-FLAG at 0, 4 and 8 days after initiation of decidualisation with cAMP and MPA showing gain of an epithelioid phenotype. The scale bars are 400 μm.

